# Multi-seasonal eDNA metabarcoding highlights a resurgence in fish diversity across a severely impacted estuarine ecosystem

**DOI:** 10.1101/2025.07.09.663915

**Authors:** Jake M. Jackman, Naiara Guimarães Sales, Chiara Benvenuto, Andrea Drewitt, Andrew Wolfenden, Peter E. Robins, Ilaria Coscia, Allan D. McDevitt

## Abstract

Aquatic ecosystems have been in an alarming state of decline for decades. Therefore, there is an urgent need for more accessible and rapid monitoring methods that will ultimately allow for restoration programmes to be implemented. In this study, we deployed environmental DNA (eDNA) metabarcoding as a fish monitoring tool in the Mersey estuarine system (UK), where ecosystem productivity has been severely impacted by water quality degradation since the Industrial Revolution. Monthly water samples were collected over a year (2023-2024) throughout the estuary, covering saline, brackish and freshwater zones. Overall, 69 species of fish were detected, increasing the number of known species within the estuary in comparison to both historical and contemporary records (46 and 39, respectively), with a generally higher observed richness within the upper estuary sampled zones where the water chemistry is predominantly freshwater. Notably, we identified several species returning to the system for the first time since pre-industrial times. Peak species richness was recorded during the winter season (December-February). Species compositions varied significantly by month and spatially by zone within months, but not when grouped seasonally. Furthermore, around ∼15% of the species detected were diadromous, with the endangered Atlantic salmon *Salmo salar*, for example, being frequently detected during its key migratory period. This study highlights the resurgence in fish diversity in a once biologically depleted estuarine ecosystem and demonstrates how eDNA metabarcoding can be implemented for detecting historically absent species, thereby enhancing restoration monitoring efforts globally.

## Introduction

The world’s aquatic ecosystems are in a perpetual state of pressure from anthropogenic stressors (Dubois et al., 2018), with freshwater and estuarine systems being heavily impacted (Kennish, 2002; Mahoney & Bishop, 2017; Dudgeon & Strayer, 2025). The deterioration of these habitats is driven by factors such as pollution, overfishing, the introduction of invasive species, habitat loss, and is augmented by climate change (Cai et al., 2021). The heterogeneous chemistry of estuarine waters, formed through the mixing of freshwater and terrestrial loads from rivers with coastal saltwater, creates a unique range of habitats that support a diverse array of species (Ray, 2005). These conditions are especially beneficial for the early life stages of many marine organisms (Tagliapietra et al., 2009). Beyond supporting early life stages, estuaries are also crucial for the migratory behaviour of fishes (Potter et al., 2015). Migratory patterns, along with other behaviours such as spawning and foraging, are integral for understanding ecosystem health, as they provide insights into how species interact with and adapt to their changing environment (Franco et al., 2008; Whitfield et al., 2023). For many diadromous species, coastal ecosystems serve as key stopover points where individuals rest, feed, and prepare for migration, playing an integral role in their life cycle (Chalifour et al., 2019). Between 1970 and 2016, populations of migratory freshwater fish have plummeted by an average of 76%, with the most pronounced losses observed in Europe (Living Planet Index for Migratory Freshwater Fish, 2020).

Monitoring fishes in complex ecosystems, such as estuaries and rivers, can be challenging, expensive, and potentially yield inconsistent results depending on the method used (Fullerton et al., 2010; Elliott et al., 2022). Environmental DNA (eDNA) metabarcoding has emerged as an established and effective technique for monitoring fish biodiversity globally (Garlapati et al., 2019; Pawlowski et al., 2020; Schenekar, 2023), with the advantages of being non-invasive and providing either detections comparable to, or superseding, more traditional fish monitoring methods (McDevitt et al., 2019; McElroy et al., 2020). There is a plethora of eDNA studies demonstrating the ability and potential of eDNA metabarcoding for the effective monitoring of fishes across both marine and freshwater ecosystems (e.g., Stat et al., 2017; Bessey et al., 2020), and increasingly in estuarine ecosystems also (Jackman et al., 2024; Cunningham et al., 2024). However, few studies have attempted to capture full-season cycles of fish diversity using eDNA metabarcoding in estuarine systems (Stoeckle et al., 2017; Gibson et al., 2024; Cunnington et al., 2024). Seasonality, driven by factors such as temperature, rainfall, and day length, plays a crucial role in estuarine ecosystems. These periodic shifts can influence breeding cycles, food availability, and habitat conditions (Arevalo et al., 2023). The dynamic nature of estuaries further shapes these processes: salinity fluctuates hourly with tides, episodically with rainfall, and seasonally with evaporation and heating (Bertram et al., 2021). Such variability makes migration an essential but challenging behavioural adaptation for many species.

This study investigates fish biodiversity in the Mersey Estuary (UK; Fig. 1), an example of an estuarine ecosystem that has suffered from long-term and lingering anthropogenic degradation since the Industrial Revolution (Lallias et al., 2015). It is estimated that prior to the Industrial Revolution, the Mersey was home to a wide variety of fish species, with the estuary playing an important role in supporting both freshwater and marine species, including migratory species like Atlantic salmon *Salmo salar*, European smelt *Osmerus eperlanus,* and European eel *Anguilla anguilla* (Mersey Rivers Trust, 2021). Post-Industrial Revolution, however, the Mersey was considered biologically dead up until ∼50 years ago (Jones, 2000, 2006). However, the implementation of new water quality legislation in the 1980s through the efforts of the Mersey Basin Campaign (Alexander & Harper 1989) aimed to improve the water quality and environmental conditions of the Mersey and its surrounding area. The effort focused on reducing industrial pollution, improving sewage treatment and wastewater management, restoring river habitats, enhancing biodiversity, and raising public awareness about environmental issues. As a result of these efforts, significant improvements in the overall health of the Mersey estuary were achieved.

**Figure 1.**
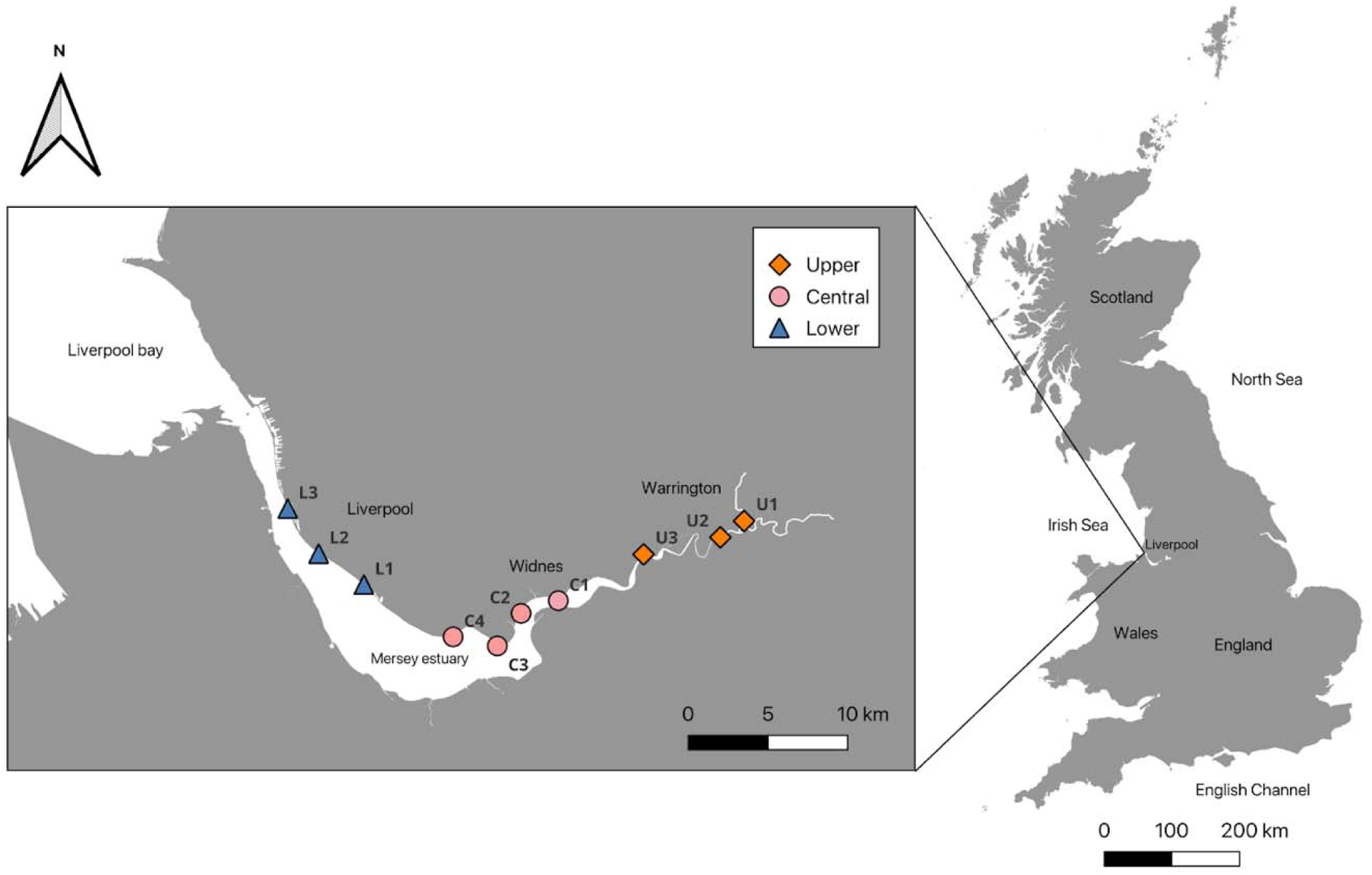
Sampling sites on the Mersey Estuary: three in the upper zone (orange diamonds), four in the central zone (pink circles) and three in the lower zone (blue triangles).

Before widespread industrialisation throughout the system, around 46 species were estimated to occur in the Mersey, and between 2015 and 2020, 39 species were documented as having returned (see Table A3 for full species lists; The Mersey Rivers Trust, 2021). Among the 46 pre-industrial species, several are diadromous (∼17%), relying on the estuary as a crucial migratory route. In the more recent 2015–2020 survey (39 species), around ∼25% of the returning species were diadromous, indicating some recovery of ecological connectivity within the estuary. However, the estimated abundances of these migratory species are likely still far from pre-industrial levels, especially considering their disrupted migration routes and altered environmental conditions. While their return is promising, the continued fragility of these populations underscores the ongoing challenges in restoring full ecological health to the system. This means that monitoring programmes would need to take this into account and limit invasive approaches and general handling of the specimens. For this reason, eDNA metabarcoding offers the potential for applying a non-invasive method to obtain estimates of biodiversity in the system.

This study aims to apply eDNA metabarcoding as a fish monitoring method across large spatial and temporal scales in the Mersey Estuary across a marine to freshwater gradient, collecting samples during a 12-month period using an optimised eDNA protocol (Jackman et al., 2024). In doing so, we aim to detect broadscale fish diversity, compare seasonal differences in richness estimates, and highlight the presence/absence of key diadromous species (UK Biodiversity Action Plan (BAP) species; Joint Nature Conservation Committee, 2007). The findings will reinforce the prospect of eDNA metabarcoding as a routine biomonitoring method in similar systems worldwide and help inform the many organisations currently working to preserve and increase fish biodiversity within this ecosystem.

## Methods

For this study, 10 sampling locations were selected (Jackman et al., 2024; Fig. 1) on the northern banks of the Mersey Estuary. Three sites were sampled in the upper freshwater zone (U1, U2 and U3) and three in the lower marine zone (L1, L2 and L3). As the central zone is the largest area of the entire system (∼5 km in width) and features added complexity due to the convergence of freshwater and marine water, four sites (C1, C2, C3 and C4) were selected here.

### Sample collection

Samples were collected during 12 consecutive months from September 2023 to August 2024. Samples were collected during three subsequent days in each month (Table A1), always during high tide due to multiple sampling locations being frequently dry at low tide and to harmonise the hydrodynamic conditions at the time of sampling, as much as possible (Jackman et al., 2024). Five water sample replicates were taken at each of the ten sites using 0.45 µm Sterivex filters (Merck Millipore), following an optimised protocol comparing different filtering methods (Jackman et al., 2024). Water samples were collected in individual 1L sterilised buckets, using one bucket per replicate, and filtered using the Sterivex 0.45 µm filters. A single 1L field blank water replicate was taken at the beginning of each sampling day using sealed bottled water. Each filter used a 60 ml syringe to manually pass the water through the filter membrane. Filtration of the water samples was performed on-site immediately following collection. Once each filter replicate became clogged, volume filtered was recorded (Table A2), and the filter was sealed in an individual sterile air-tight bag and stored at −20°C in the laboratory until processed. In total, 636 samples were collected across the study, including 600 eDNA samples (10 sites x 12 months x 5 replicates) and 36 field negative controls (3 days x 12 months).

### DNA extractions, PCR amplification and Sequencing

All DNA extractions were performed in a dedicated eDNA clean room at the University of Salford, decontaminated via UV for at least 6 hours before extractions; all personnel wore full PPE. The DNA extractions followed the Mu-DNA protocol (Sellers et al., 2018) tailored for water samples (see Jackman et al., 2024 for further details). During each extraction session, an extraction negative control (extraction reagents excluding the DNA template) was included to account for possible contamination.

DNA was PCR amplified using the fish-specific Tele02 primers (forward: 5′-AAACTCGTGCCAGCCACC-3′; reverse: 5′-GGGTATCTAATCCCAGTTTG-3′; Taberlet et al., 2018) according to the protocol described in Jackman et al. (2024). Both positive (*Hoplias malabaricus,* a neotropical freshwater species absent in the UK) and negative (nuclease-free water) PCR controls were included to account for possible tag jumping and contamination. Due to a high number of samples (600 eDNA samples, 36 field blanks, 24 extraction blanks, 16 PCR blanks and 16 positive controls), PCRs were prepared across eight libraries as follows: libraries 1-4 each contained 75 eDNA samples, four field blanks, three extraction blanks, two positive controls and two PCR negative controls (86 samples in total). Libraries 5-8 each contained 75 eDNA samples, five field blanks, three extraction blanks, two positive controls and two PCR negative controls (87 samples in total). PCRs were performed in triplicate for each sample. Each sample was amplified using a unique 8 bp oligo-tag attached to the forward and reverse primers and a variable number (2-4) of leading Ns (fully degenerate positions) to increase variability in amplicon sequences. PCR amplification was conducted using a single-step protocol to minimize bias in individual reactions. The PCR reaction consisted of a total volume of 20 µL including 10 µl AmpliTaq Gold™ 360 Master Mix (1X; Applied Biosystems); 0.16 µl of BSA (5mg); 1 µl of each of the two primers (5 µM); 5.84 µl of ultra-pure water and 2 µl of eDNA template. All PCR amplifications of libraries were performed under the following thermocycling conditions: 95°C for 10 min, followed by 40 cycles of 95°C for 30 s, 60°C for 45 s, and 72°C for 30 s, and a final elongation of 72°C for 5 min.

Replicates were then pooled, and samples were visualised on a 1.2% agarose gel stained with GelRed® to check for successful amplification of target fragments. PCR products were then purified with HighPrep™ PCR Clean-up System magnetic beads using a 1:1.1 ratio for a left-sided size selection. The purified libraries were visualised on the Agilent 2200 TapeStation using High Sensitivity D1000 ScreenTape (Agilent Technologies). This indicated secondary non-target products on the right side of the target fragment, which was removed by right-sided size selection (1:0.8 ratio for all libraries). Size-selected DNA was quantified using a Qubit™ 4.0 fluorometer with the Qubit™ dsDNA HS Assay Kit (Invitrogen). Based on the total DNA concentration, each library was diluted to 20ng/ul at a volume of 50ul for library preparation. End repair, Adapter ligation and library PCR amplification were performed using the KAPA HyperPrep Kit according to the manufacturer’s protocol. Libraries were quantified using quantitative PCR (qPCR) on a MIC qPCR system (Bio Molecular Systems) with the NEBNext® Library Quant Kit for Illumina® (New England Biolabs). All prepared libraries were sequenced using the Illumina® NovaSeq X Plus PE150 (Novogene, UK) with a minimum of 6 GB of raw data generated per library.

### Bioinformatic analysis

The bioinformatics analysis was completed using the OBITools metabarcoding software 1.2.11 (Boyer et al., 2016). Read quality was checked with *fastqc,* and low-quality ends were trimmed for downstream analysis. We used *illuminapairedend* to merge all paired reads showing a quality score >30 and *ngsfilter* to demultiplex samples based on their unique barcodes. Sequences were filtered via *obigrep* to remove singletons and reads out of the expected length range (129-209 bp) and dereplicated via *obiuniq*. We removed chimeras with *uchime* (Edgar et al., 2011) and clustered the remaining sequences into Molecular Operational Taxonomic Units (MOTU) with *swarm* (Mahé et al., 2015), setting the threshold to d = 1. Sequences were assigned taxonomy information using a DNA reference library dataset for fish species of the United Kingdom (UK), derived from the NCBI GenBank and Barcode of Life BOLD databases (Meta-Fish-Lib v255). The reference dataset includes species for both freshwater and marine UK species (Collins et al., 2021). Due to limitations in the reference database, the final dataset had to be manually curated using the NCBI nucleotide database (Altschul et al., 1990), to improve taxonomic assignments by checking MOTUs identified at the genus level (see Jackman et al., 2024 for further details). All subsequent analyses were performed in Rv4.3.1 (R core team 2023).

MOTUs/reads originating from sequencing errors or contamination were removed, as well as non-fish and non-target species (e.g., human and domestic species reads). To address MOTUs that were containing potential contaminants, the maximum number of reads recorded in the controls (field collection blanks, DNA extraction blanks and PCR blanks) were removed from all samples. Finally, all MOTUs with < 5 reads were removed from the final dataset. The final taxonomic assignment was conducted according to current fixed general thresholds: MOTUs were assigned at the species level when matching the reference sequence with > 98% in line with other UK fish studies (Hallam et al., 2021; Rourke et al., 2022). After all filtering stages, all libraries were separated in their respective sampling months (September 2023 - August 2024).

### Data analysis

Seasonality of the data was determined based on the typical seasonal cycles in the United Kingdom, following the classification by the UK Met Office: autumn (September, October, and November), winter (December, January, and February), spring (March, April, and May), and summer (June, July, and August) (<u>UK Met Office</u>, 2024). These periods are also aligned with key migratory patterns of estuarine fish species, such as autumn migrations to overwintering sites (Graham & Harrod, 2009) and spring migrations associated with spawning, as observed in several species across UK estuaries (Ibbotson et al., 2013).

To evaluate the relationship between sequencing depth and biodiversity, we conducted a Kendall’s rank correlation test (Kendall’s Tau) between the total read count (sequencing depth) and total observed species richness across all samples. This non-parametric correlation analysis was performed using the cor.test() function v4.3.1 (stats package; R core Team, 2023), chosen due to the non-normality of both variables as confirmed by Shapiro-Wilk tests (Total reads: W = 0.69266, p < 0.001; Total richness: W = 0.94934, p < 0.001). The analysis included all 12 sample months to assess the global relationship independent of temporal or spatial factors.

The number of species detected through the eDNA survey was compared with the species listed in the Mersey Rivers Trust (2021) report, which details species estimated to inhabit the Mersey pre-industrial revolution and species detected during a five-year survey between 2015-2020. This comparison was performed using a Venn diagram (VennDiagram v1.7.3; Chen & Boutros, 2011). The brook lamprey *Lampetra planeri*, European river lamprey *Lampetra fluviatilis*, and mullet species, the thick-lipped grey mullet *Chelon labrosus* and thin-lipped mullet *Chelon ramada*, were excluded from this comparison because each species pair is indistinguishable using the target 12S rRNA region. Since we could not determine whether all or only one species from each genus was detected, we could not conclusively state whether they remain exclusive to the pre-industrial list or were also present in the eDNA list.

Species richness across the ten sampled sites and seasons was visualised using boxplots (generated with ggplot2 v3.1.5; Wickham, 2016). Differences in richness between seasons were assessed using the Kruskal-Wallis test, followed by Dunn’s post hoc test for pairwise comparisons. Each species from the eDNA survey and the 2021 report has been classified to a physiological saline tolerance (F = freshwater, M = marine, and B = brackish) using classifications as described per species in FishBase (FishBase, 2025; Table A3).

To assess the overlap in species detected between sampling months and seasons, UpSet plots were created using the **“**UpSetR**”** package v1.4.0 (Conway et al., 2017). UpSet plots provide an efficient visualisation of intersections (number of species) between sets (sample sizes), making them particularly useful for identifying shared and unique species across different sampling periods. To assess temporal changes in species composition, analyses were first conducted at the monthly scale before examining broader patterns at the seasonal level using non-metric multidimensional scaling (NMDS). A PERMANOVA was performed using the Bray-Curtis dissimilarity index to test for statistical differences in species composition across the 12 sampling months and the four seasons, and was applied to test for spatial variation between zones within each month. This approach first allowed for an assessment of fine-scale temporal variation, capturing potential short-term shifts in community structure which may not be visible on a broader, seasonal scale. NMDS ordinations were generated using the “vegan” package v2.6.8 (Oksanen et al., 2022) to visualise patterns of similarity and dissimilarity across months and seasons. This is followed up by a pairwise comparison, conducted using the pairwiseAdonis function to assess the differences in species composition between each pair of months and seasons.

Finally, notable migratory species (e.g., protected UK BAP fish species), the season and site they were detected have been visualised using an alluvial plot, using the package “ggalluvial” (Brunsen, 2023). For this comparison, *Lampetra* spp. was included in the analysis as *L. fluviatilis* is a key migratory species (but with the caveat that this could be *L. planeri*; see above). This was also performed for all detected species classified as freshwater/brackish or marine/brackish (Table A3), showcasing their physiological saline tolerance and the estuarine habitat zone in which they were detected.

## Results

The eDNA dataset obtained after quality-checking and filtering allowed for the detection of at least 69 unique fish species in total (Table A3). *Lampetra* spp. and *Chelon* spp. were retained at the genus level for subsequent analyses (see Methods). The highest number of species was detected in March 2024 (*n* = 46) and the lowest in June 2024 (*n* = 30; Fig. 2). The Kendall’s Tau correlation test revealed a weak but statistically significant positive correlation between total read count and species richness (τ = 0.34, p < 0.001; Fig. A1). However, four of the months with generally the lowest read counts returned the highest species richness values (Fig. 2).

**Figure 2.**
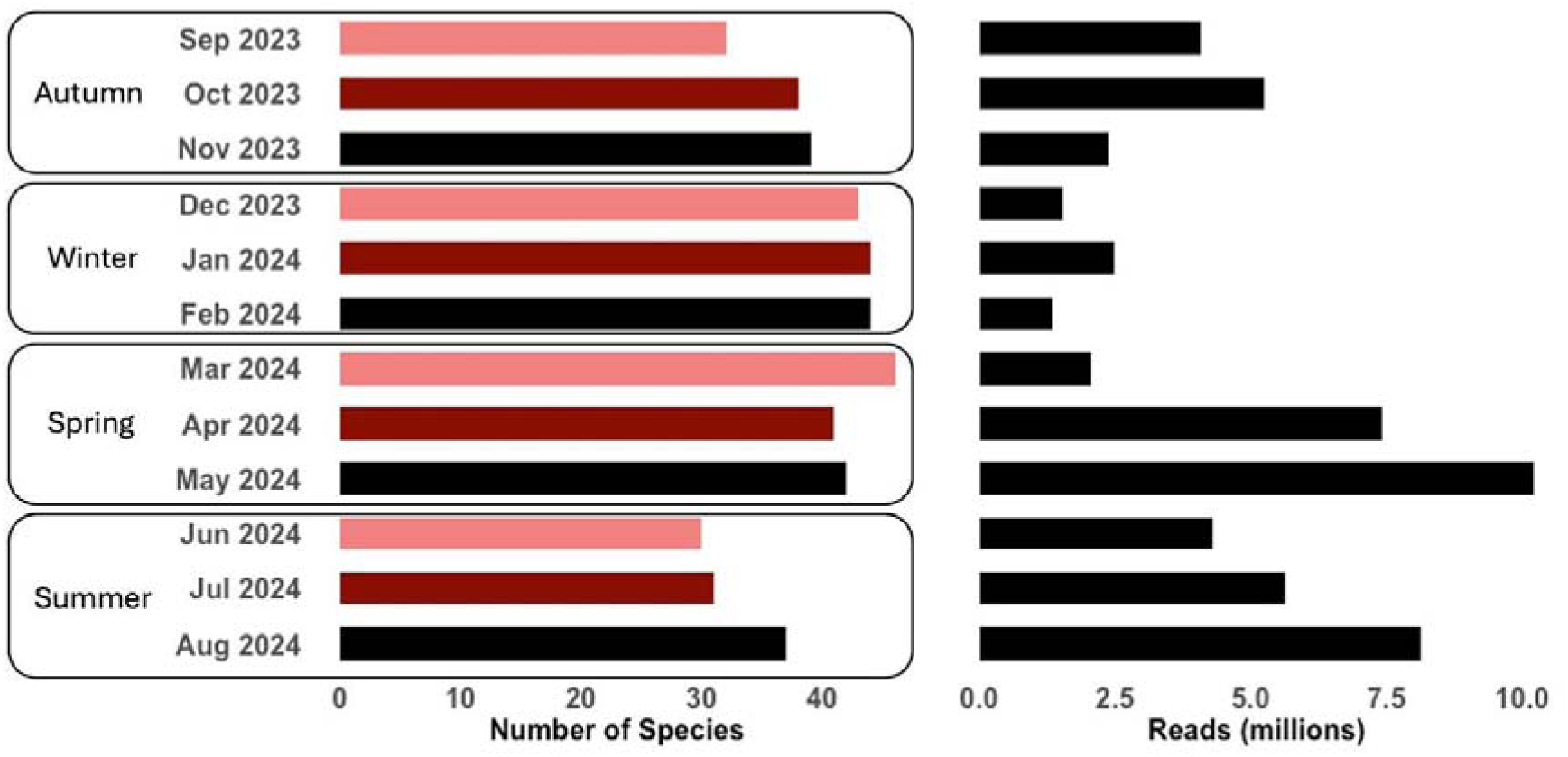
Total number of species and reads (x-axes) recovered by each of the 12 sampling months by season (y-axis).

The species lists from the Mersey Rivers Trust report (2021), alongside the species detected in the eDNA survey for this study, document a total of 81 unique species across three species lists (Table A3). A Venn diagram (Fig. 3) visualises the species overlap and exclusivity across these datasets. Of the total species, 26 species are shared among all three lists (∼34%). Additionally, 25 species were exclusively detected through the eDNA survey (32%), while four species were unique to the pre-industrial estimates (5.2%), and four species were exclusive to the 2015-2020 survey data (5.2%).

**Figure 3.**
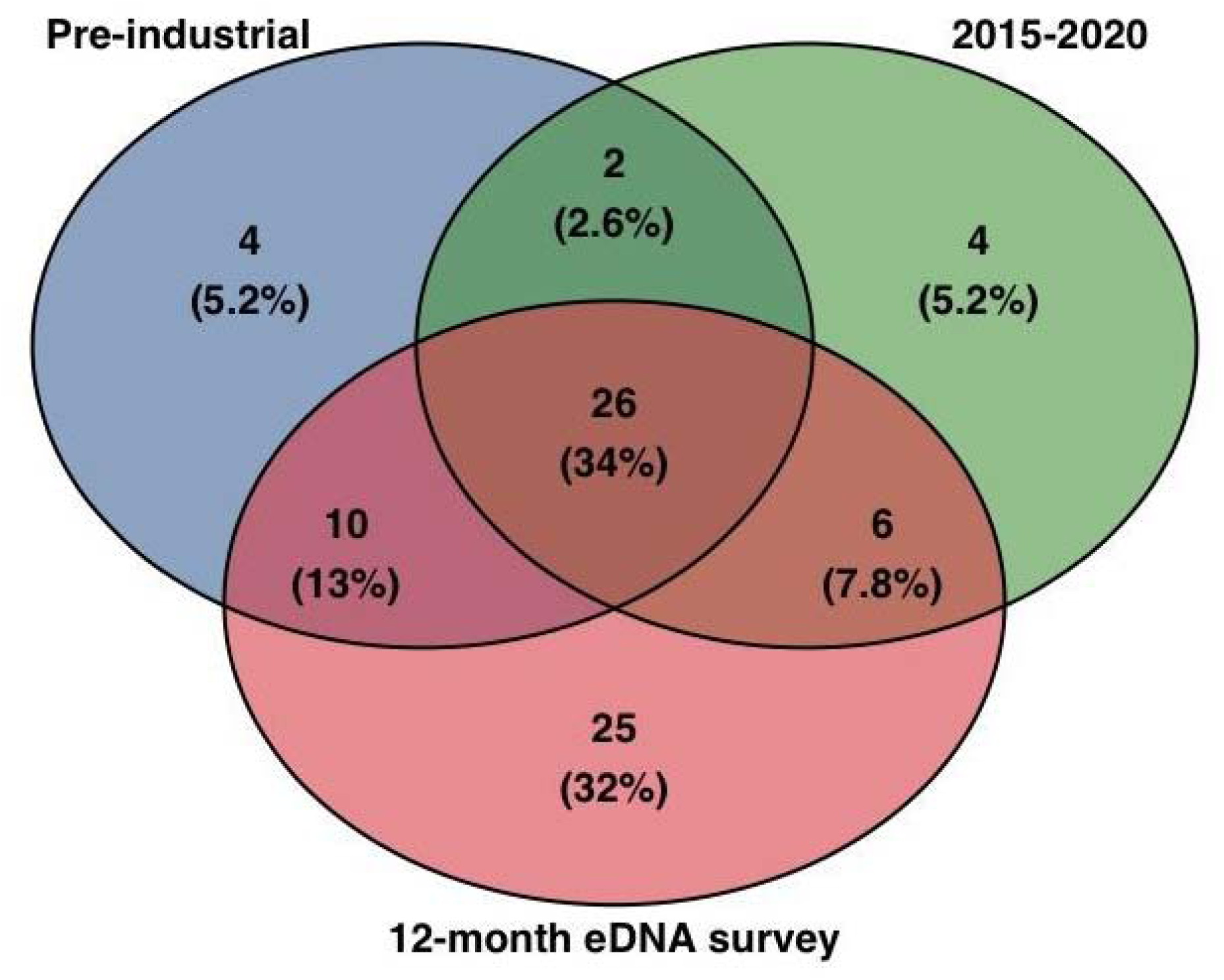
Venn diagram that depicts the total species shared and exclusively detected for each of the compared species lists.

The four species that are exclusive to the pre-industrial list are the burbot *Lota lota* (now extinct within the UK; NatureServe & Freyhof, 2024), torpedo ray *Torpedo sp.*, shore rockling *Gaidropsarus mediterraneus*, and the European sea sturgeon *Acipenser sturio* (deemed ‘possibly extinct’ in this region; IUCN, 2022). The four species that are exclusive to the 2015-2020 survey are the butterfish *Pholis gunnellus*, sea lamprey *Petromyzon marinus*, poor cod *Trisopterus minutus* and the tub gurnard *Chelidonicthys lucernus*.

Species richness variations across the sampled sites and months are evident (Fig. 4). Across all seasons, species richness is consistently higher in the Upper zone (U1–U3) compared to the Central (C1–C4) and Lower (L1–L3) zones. In spring and summer, richness gradually increases from the Lower to the Upper sites, with the highest values observed between U1 and U3. Autumn follows a similar trend, although there is more variability in richness at the Central sites. Winter shows the most pronounced separation, with the Upper sites maintaining the highest richness, while richness at Lower and Central sites remains more constrained, with lower median values and fewer outliers.

**Figure 4.**
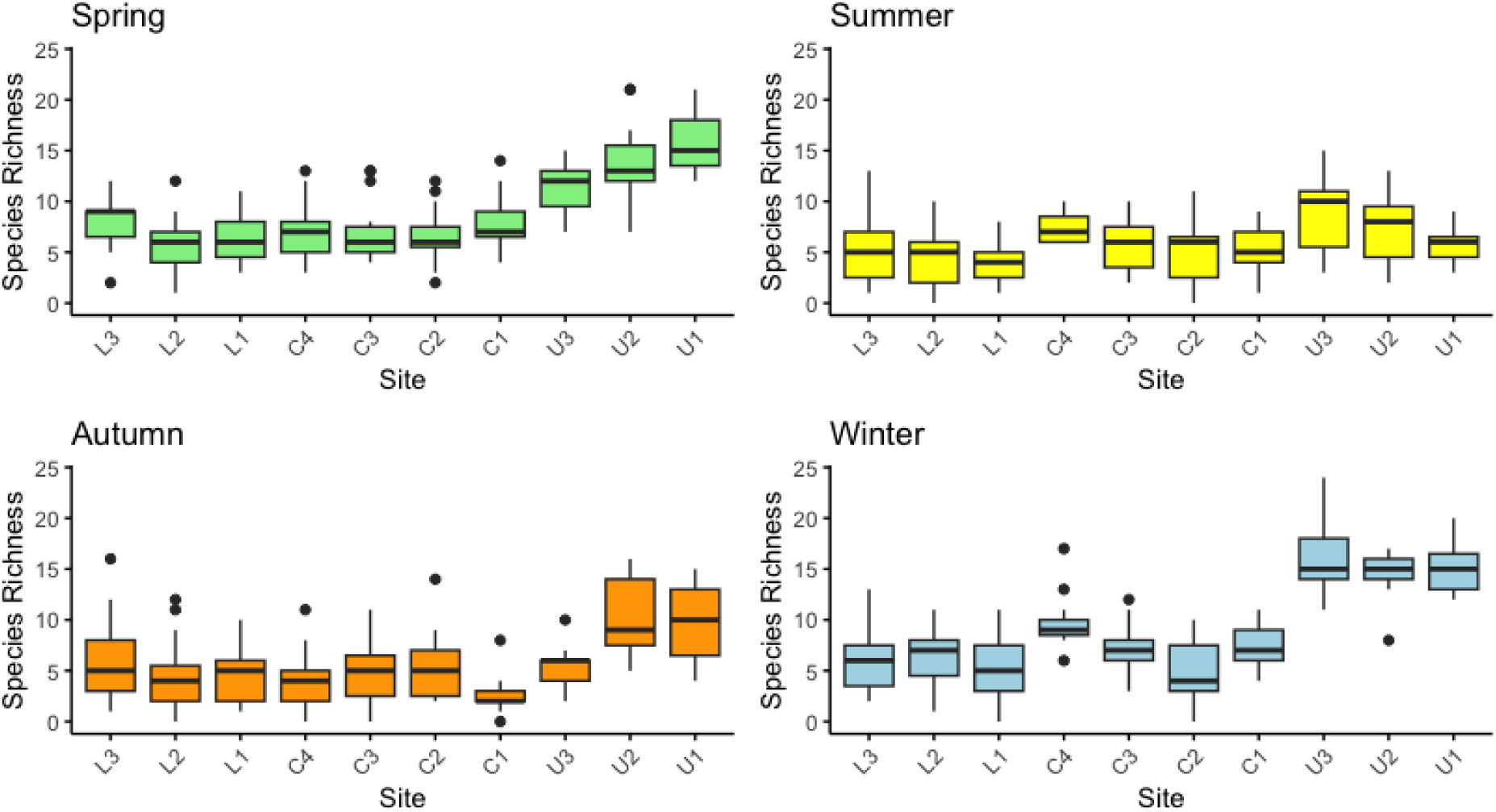
Species richness detected by each sample site for each season of the year (spring, summer, autumn and winter).

Overall, species richness varied significantly across the four seasons (χ² = 83.415, df = 3, p < 0.001; Fig. A2). Dunn’s post hoc pairwise comparisons indicate that species richness differs significantly between multiple comparisons, except for spring vs. winter and summer vs. autumn (see Table A4 for all comparisons). The lack of significant differences between spring and winter, as well as summer and autumn, suggests more stable richness levels within these seasonal pairs, whereas transitions between other seasons show stronger shifts in species richness.

Species detection varied across both individual sampling months and broader seasonal scales (Figs. 5A and B). In total, twelve species were detected exclusively within individual months (Fig. 5A), representing approximately 17.3% of the total richness. When considering seasonal patterns (Fig. 5B), the highest number of exclusive detections occurred in winter, with eight species (∼12% of the total richness). Spring had four exclusively detected species (∼5.8%), autumn had two (∼3%), and summer had the lowest, with only one species (∼1.5%) (Table A3).

**Figure 5.**
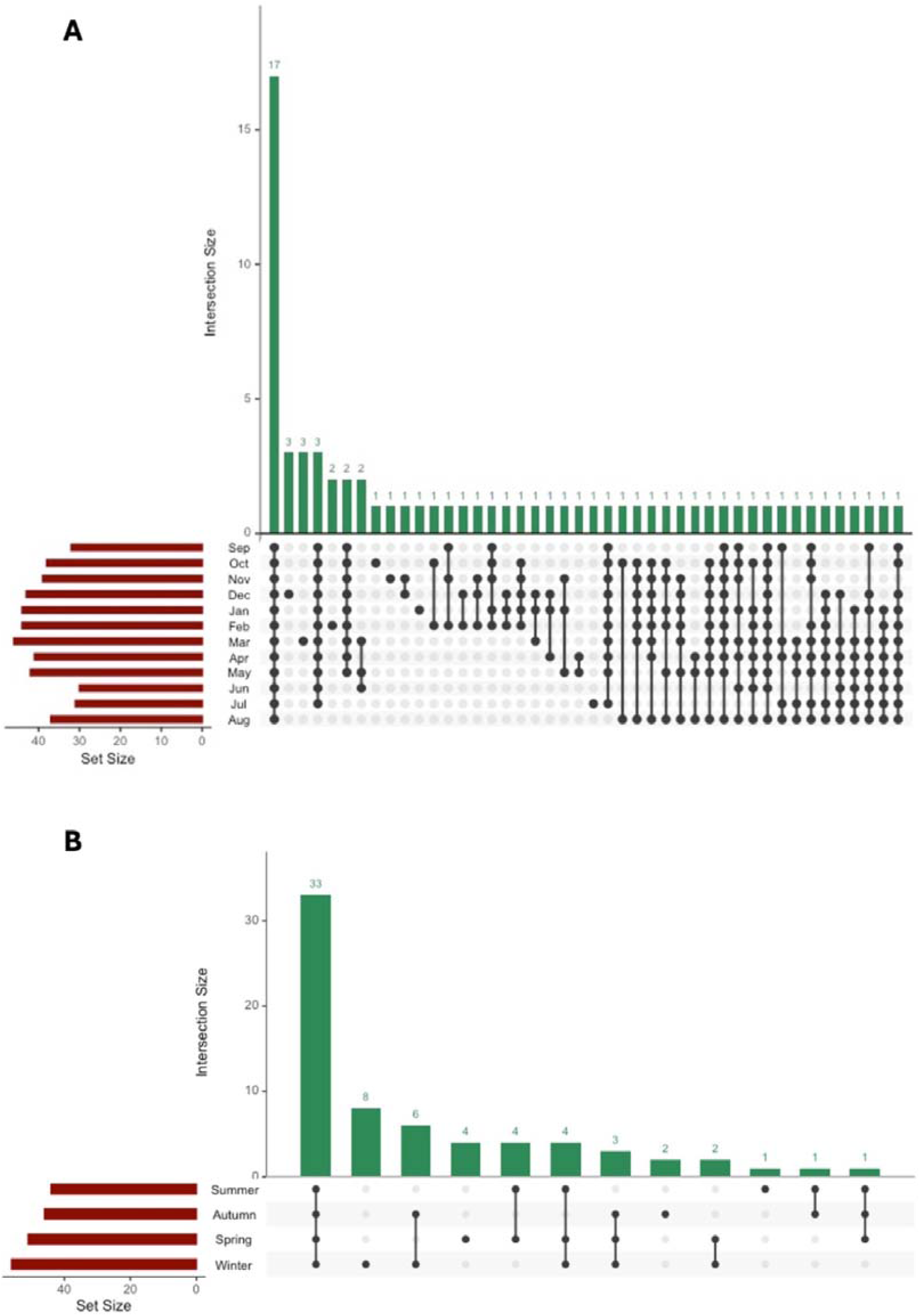
Upset plots of the number of species detected by month (set size) and exclusively between months (intersection size) (A) and the number of species detected by season (set size) and exclusively between season (intersection size) (B).

The eight species exclusively detected in winter were European hake *Merluccius merluccius*, pollack *Pollachius pollachius*, longspined bullhead *Taurulus bubalis*, tench *Tinca tinca*, Atlantic horse mackerel *Trachurus trachurus*, corkwing wrasse *Symphodus melops*, turbot *Scophthalmus maximus* and lesser pipefish *Syngnathus rostellatus.* The four in spring were Crucian carp *Carassius carassius*, crystal goby *Crystallogobius linearis*, common seasnail *Liparis liparis* and European anchovy *Engraulis encrasicolus.* The two in autumn were big-scale sand smelt *Atherina boyeri* and striped red mullet *Mullus surmuletus.* The single exclusive detection in summer was the starry smoothhound *Mustelus asterias*.

When assessing temporal species composition changes, a PERMANOVA analysis revealed that species composition varied significantly across the 12 sampling months (*F* = 5522.8; *p* = 0.001; Fig. A3) and significantly between the sampled zones within each month (Table A5). This result suggests that species assemblages shift meaningfully over time, indicating strong temporal community structuring. All post hoc pairwise comparisons between months can be seen in Table A6. When fish community composition was compared across the four seasons, it was found that differences were not significant (PERMANOVA *F* = 1.5172; *p* = 0.159; Fig. 6).

**Figure 6.**
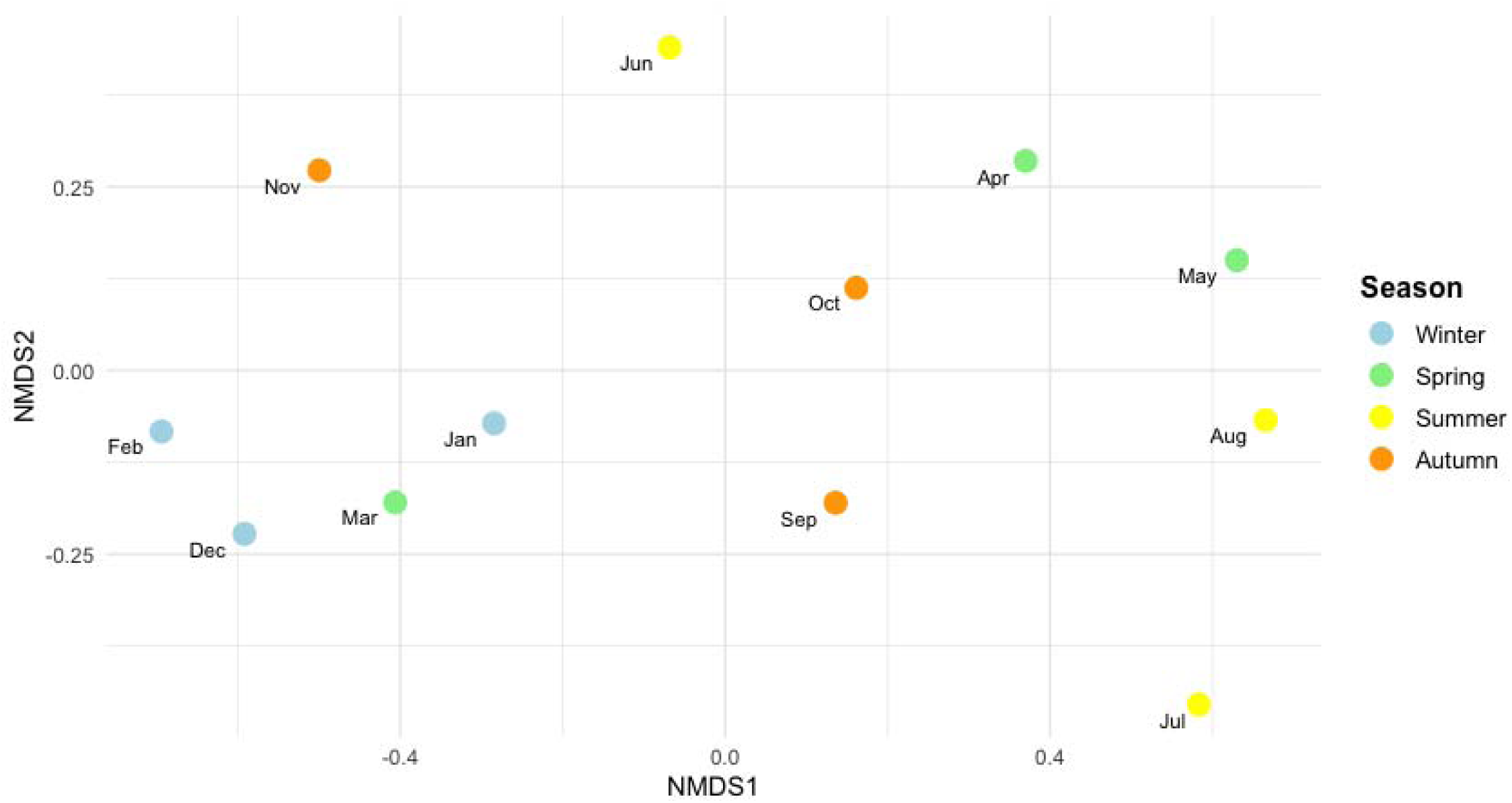
A nonmetric multidimensional scaling (NMDS) analysis was performed using Bray–Curtis dissimilarity (stress = 0.071). The plots show all eDNA samples aggregated by sampling month and coloured by different seasons.

Across the dataset, several migratory species, and listed on the UK BAP list, were detected. The European eel *A. anguilla* was present across all sampling months and seasons and at all sampling sites, and the brown trout *S. trutta* was detected in all seasons and sites (the month of July being the only exception). Atlantic salmon *S. salar* was found in winter (December, January, and February) at sites U1, U2, U3, L1, and L2, and in autumn (October and November) at sites U1, U2, and U3. Detections of *Lampetra* spp. were recorded in spring (April) at site L1 and in winter (December– January) at sites U3 and C1. European smelt *Osmerus eperlanus* occurred across all seasons at sites U1, U3, C2, C3, C4 and L3 (Fig. 7).

**Figure 7.**
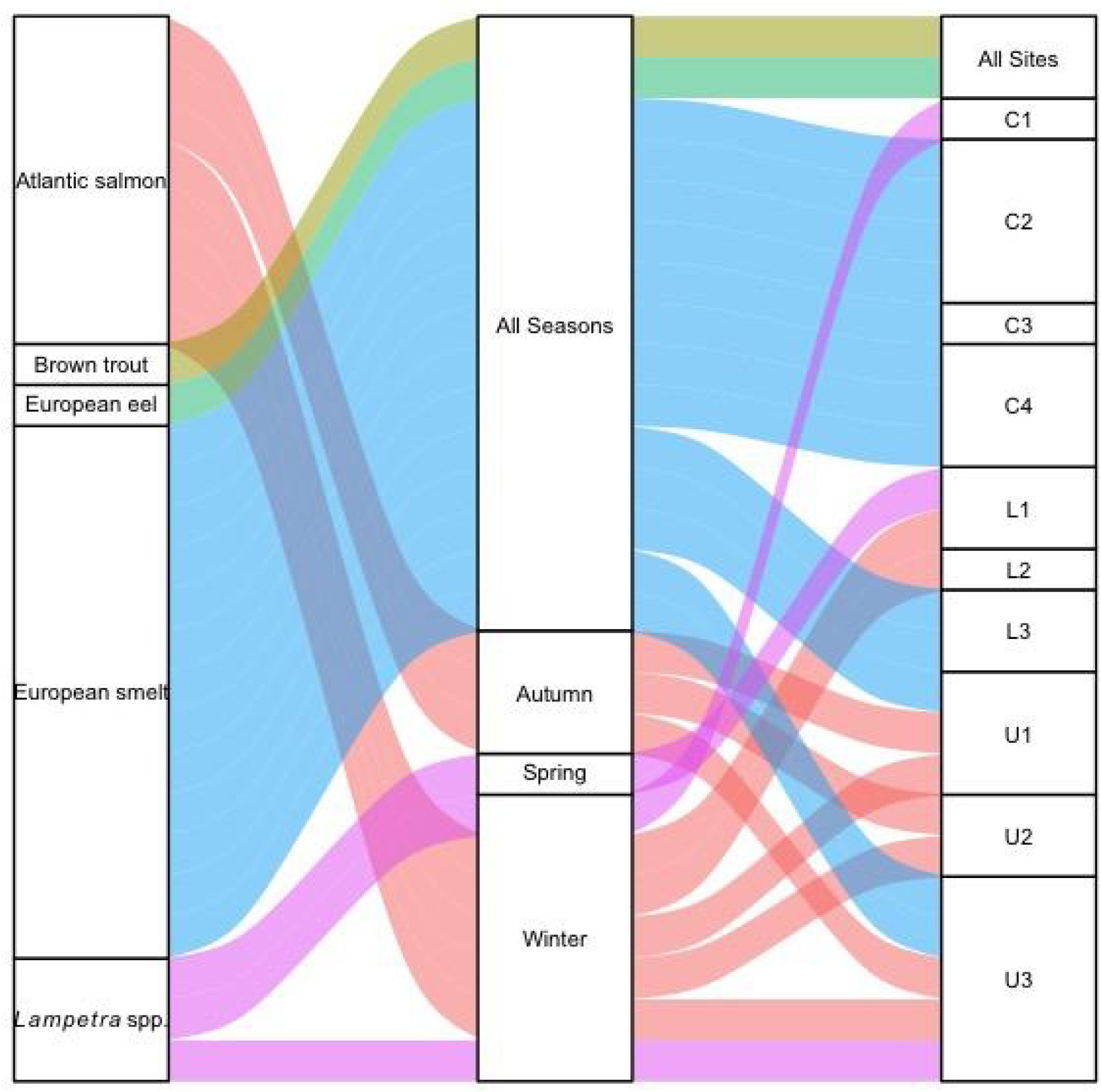
Alluvial plot depicting key migratory species detections with their associated sampling site(s).

Of the 69 detected species (Table A3), 20 are classified as freshwater/brackish tolerant and 37 are classified as marine/brackish tolerant. Of the 20 freshwater/brackish species, 31.2% were detected in the central zone and 27.1% were detected in the lower zone. Of the 37 marine/brackish species, 34.7% were detected in the central zone and 24% were detected in the upper zone (Fig. A4).

## Discussion

With climate change intensifying, estuarine ecosystems face mounting pressures from rising sea levels (projected to increase by +1m this century), more frequent storm surges, and ocean warming (with temperatures potentially rising by +4°C by 2100; IPCC, 2021). These shifts exacerbate existing anthropogenic stressors, including altered river flows, increased nutrient/sediment loads, and pollution from industrial and urban sources, legacies of the Industrial Revolution that persist in many systems (Clausen & York, 2008; Arthington et al., 2016). Such cumulative impacts have led to widespread declines in coastal marine fish populations, underscoring the need for efficient, scalable monitoring tools like eDNA metabarcoding. In this study, we assess fish biodiversity in a heavily modified post-industrial estuary. Despite historical degradation, our data reveal a rich assemblage of species across seasons, many of which have reappeared after decades of absence, a likely result of the restoration initiatives implemented in the 1980s (Kim & Batey, 2021).

### Species richness and fish diversity in the Mersey

In total, at least 69 distinct species of fish were identified across 12 months of sampling. The number of species detected here exceeded all previous estimates, with all current documented species lists recovered through more traditional methods featuring a maximum of ∼46 species recorded (Mersey Rivers Trust, 2021). All species detected through eDNA analysis were cross-examined against expected species for the geographic location and are consistent with the anticipated composition for these environments (Baldock & Dipper, 2023). At least five UK Biodiversity Action Plan (BAP) priority fish species were detected: the European eel, the brown trout, the Atlantic salmon, the European smelt and *Lampetra* spp. (Joint Nature Conservation Committee, 2007), representing an increase in species detected in comparison to previous eDNA work conducted during November 2022-January 2023 (Jackman et al., 2024). The consistent identification of UK BAP priority species further underscores the power of eDNA to detect species of conservation concern, many of which may exist in low populations or occupy complex habitats (Beng and Corlett, 2020). This is further substantiated by the detection of thick-lipped grey mullet *C. labrosus* or thin-lipped mullet *C. ramada* (designated here as *Chelon* spp. due to indistinguishable 12S rRNA gene sequences (Cunnington et al., 2024) and the transparent goby *Aphia minuta*, a set of species/genera previously assumed to be absent due to insufficient habitat quality (Mersey Rivers Trust, 2021). The Mersey Estuary’s ability to support more habitable conditions for these fish species underscores its ecological significance and highlights clear signs of ecological recovery over the past four decades (Hawkins et al., 2020).

We see a generally higher observed richness within the upper estuary sampled zones, in which the water chemistry is predominantly freshwater. This trend is present over the four seasons (Figs. 4 and A2). In highly dynamic and turbid environments such as tidally energetic estuaries, the suspended particles within a water sample can quickly obstruct the filter and inhibit the filtration process (Barnes et al., 2021; Hallam et al., 2021). However, water volume filtered is likely not the sole contributing factor toward the increased richness estimates (Jackman et al., 2024). Ecological factors, such as habitat structure, salinity gradients, and connectivity between saline gradient zones, can significantly influence species richness (Leibold et al., 2004; Lin et al., 2024). Furthermore, only a weak correlation between species richness and sequencing depth was observed (Fig. A1), suggesting that other biological and ecological factors may drive richness variation, and the balanced library preparation ensures that differences in sequencing depth are unlikely to bias richness estimates between the sampled zones. In addition, the classification of species (Table A3) to either freshwater or marine indicates more marine species present in the data, and the alluvial plot, showcases prevalent abiotic influence of potential eDNA transport from the marine zone to the upper (25% of marine species detected in the upper freshwater zone; Fig. A4) contributing to the elevated richness observed in this zone.

The observed richness is notably higher during the winter and spring seasons, which may provide valuable insights into the ecological dynamics influencing eDNA persistence in the water column. While some winter-exclusively detected species (e.g., European hake, pollack, and turbot) are typically associated with offshore movements in winter (Casey & Pereiro, 1995; Imsland et al., 1996; Gonse et al., 2025), their detection in the system during this season contrasts with migratory behavioural expectations. Similarly, year-round residents like tench and long-spined bullhead were unexpectedly detected only in winter, further complicating the identification of clear behavioural patterns underlying these observations. The Spring season is a critical time for fish spawning (Wright & Trippel, 2009), which could further explain the elevated richness and DNA reads observed during this period. Spawning represents a biologically intense phase for many fish species, during which large quantities of genetic material, such as gametes, are released into the water (Alix et al., 2020). Our analysis of seasonal reads reveals a higher abundance of eDNA reads in the spring (Fig. 2), suggesting that the heightened eDNA concentrations and observed richness during spring may be driven by the reproductive behaviours of fish, supporting studies showing that spawning events contribute to the higher genetic material present in the water (Schreck et al., 2002; Tillotson et al., 2018; Alix et al., 2020; Collins et al., 2022). However, it is also worth noting that eDNA follows a state of exponential decay upon release (Sassoubre et al., 2016), so one potential explanation for this seasonal increase in observed richness in winter and spring may be the conditions associated with these seasons (less UV exposure and more temperate water conditions) contributing to the extended survival of DNA fragments in the water (Barnes et al., 2014; Strickler et al., 2015; McCartin et al., 2022), resulting in a larger window for capturing species’ DNA between shedding and filtration.

In estuarine systems, tidal patterns and the flow of water directly influence the transport of eDNA particles (Andruszkiewicz et al., 2019; Pont, 2024), which, in turn, should be considered when interpreting the results. For example, here, there are frequent instances of species detected outside of their physiological tolerances (Table A3; Fig. A4). Common marine/brackish species such as Atlantic herring *Clupea harengus*, and European sprat *Sprattus sprattus* have been detected across multiple freshwater sites and sampling months. Furthermore, we also see the opposite pattern, with freshwater/brackish species such as the common roach *Rutilus rutilus* and common bream *Abramis brama* frequently detected within the lower marine zones of the system. The strong tidal dynamics of the Mersey estuary, with its pronounced tidal asymmetry throughout its length (flood tides fill the upper estuary in as little as ∼ 2 hours with ebb tides retreating for up to ∼10 hours), extensive tidal excursion range (particularly at the estuary mouth as it flows into Liverpool Bay) and complex hydrology (The Mersey Gateway Project, 2021), can facilitate long-distance transport of eDNA. It has been documented that eDNA can be transported over large spatial scales (Fremier et al., 2019; Wood et al., 2021), with DNA from freshwater species being transported through estuarine regions to saltwater zones, and vice versa. Therefore, it is important to note that we are detecting the species’ DNA at a particular location and not necessarily the species itself. While this might be less important for monitoring an overall system in terms of species diversity/composition, as we have undertaken here, it could be critical for identifying important habitats for species of concern within these systems. This emphasises an urgent need for more integrative approaches, such as coupling eDNA surveys with hydrodynamic modelling and particle tracking methods to simulate hydrological patterns and the subsequent dispersal of eDNA (Fukaya et al., 2021; Pastor Rollan et al., 2024). Such approaches would help clarify how tidal regimes and estuarine hydrodynamics influence eDNA transport, and how these factors relate to the presence of a particular species both spatially and temporally.

### Seasonal compositions and diadromous species detections

Analysing the data on a finer monthly scale revealed significant fluctuations in species compositions (Fig. A3), indicating that short-term temporal dynamics play a crucial role in shaping the observed fish biodiversity within the Mersey. These findings reinforce the importance of implementing a robust temporal sampling strategy to accurately capture broadscale biodiversity estimates (Beentjes et al., 2019; Seymour et al., 2021). The monthly variability observed here may be driven by factors such as seasonal migrations, reproductive cycles, hydrological conditions, and environmental fluctuations that influence species presence and detectability (De Souza et al., 2016; Buxton et al., 2017). However, when the data is aggregated into broader seasonal periods, species compositions exhibit relatively non-significant variation across seasons (Fig. 6). Although several eDNA studies in estuaries have identified seasonal differences in fish community compositions (e.g., Stoeckle et al., 2017; Cunnington et al., 2024), other studies have found this not to be the case (e.g., Hallam et al., 2021; Gibson et al., 2023). Here, this consistency across seasons suggests that while some species exhibit strong temporal shifts at a more fine-scale monthly level, the overall assemblage remains relatively stable when viewed on a seasonal scale. A key factor contributing to this stability could be the presence of approximately half of all recorded species detected through all seasons (Fig. 5B). Consequently, while seasonal sampling can provide a general overview of biodiversity trends, higher-resolution monthly data (or if feasible, including more fine-scale sampling to account for tidal variations and its influence on eDNA transport) is important for detecting fine-scale temporal changes and ensuring comprehensive assessments of community dynamics.

The year-round species detected could represent resident populations or species that are less affected by short-term environmental fluctuations. However, the presence of artificial ecological barriers, such as weirs, docks and shipping canals, which are prevalent features throughout the length of the Mersey estuary (Cunnington et al., 2024), can also influence species distributions by impeding the natural migratory routes of certain fish species (Bunt et al., 2022; Rahel & McLaughlin, 2018). For instance, species that typically migrate to and from the estuary may become trapped, and therefore unable to complete their usual migratory journeys (Brönmark et al., 2014). This phenomenon could lead to the unintended permanent residency of these species within the system, altering their traditional migratory patterns (Chapman et al., 2012). The year-round presence of typically migratory species in the eDNA data, such as the European flounder *Platichthys flesus* (which migrates to deeper waters during winter; Tesch & Thorpe, 2003), may be a potential example of this, though alternative explanations related to eDNA ecology must also be considered. For instance, DNA shed by migrating individuals may persist in sediments and later resuspend into the water column, leading to detections even in the absence of live fish (Jerde et al., 2016; Ellegaard et al., 2020). While this study used eDNA metabarcoding, future work incorporating eRNA assays, which degrade rapidly and could potentially be more indicative of live organisms, may help distinguish active populations from residual genetic material more effectively (Littlefair et al., 2022; Janik-Superson et al., 2025). Such approaches would clarify whether these patterns reflect true behavioural shifts (e.g., halted migration or trapping) or the limitations of eDNA-based detections in dynamic ecosystems.

Overall, around ∼15% of the species detected with eDNA here are diadromous, highlighting the estuary’s role as a crucial corridor and habitat for migratory species. Atlantic salmon, for instance, rely on clean, well-oxygenated freshwater habitats for spawning, making their presence a valuable indicator of improving water quality (Thorstad et al., 2008; Smialek et al., 2021). Interestingly, the detections of salmon align with their key migratory period in late autumn to winter (Fig. 7), reinforcing the idea that conditions are becoming more suitable for their life cycle after a lengthy absence (Jones, 2006). Similarly, European smelt also depend on clean, fast-flowing water for spawning (Aarts & Nienhuis, 2003). While their presence is more sporadic, any detection suggests that improving habitat conditions may be supporting their return. The detection of *Lampetra* spp. also aligns with the spawning season of the migratory *L. fluviatilis* between April and May (Silva et al., 2015), and they also rely on clean freshwater for spawning. It is important to reiterate, however, that the 12S sequence fragment used here can not distinguish between the migratory *L. fluviatilis* and the non-migratory *L. planeri,* so a more targeted species-specific eDNA approach would be warranted for the future detection and monitoring of these two species. European eel *Anguilla anguilla*, a critically endangered species (IUCN, 2024), was frequently detected, corroborating their known presence in the Mersey over the past decade (e.g., *BBC News*, 2023; *Springwatch*, 2016). Their persistence highlights the estuary’s role in supporting at-risk diadromous species, with eDNA emerging as a promising tool for monitoring their recovery (Hillsdon, 2023; Rodriguez, 2023). Collectively, these findings signal ecological restoration in this previously degraded estuarine ecosystem, particularly for species with stringent habitat requirements (Hawkins et al., 2020).

It is also worth noting that non-target eDNA detections are present within the dataset. For this study, we used the Tele02 primer set, which is designed for the detection of teleost fishes (Taberlet et al., 2018), but we also detected two elasmobranch species in the estuary, the starry smooth-hound *Mustelus asterias* (class Elasmobranchii, infraclass Selachii) and the thornback ray *Raja clavata* (class Elasmobranchii, infraclass Batoidea). This highlights a major benefit to the eDNA as a monitoring method overall, demonstrating that non-target species can be detected as ‘molecular bycatch’ (Mariani et al., 2021). These infrequent non-target detections could be improved by re-analysing the samples using different primer sets designed to target elasmobranch species (e.g., Elas02; Taberlet et al., 2018).

### Conclusions

This study demonstrates the utility of eDNA metabarcoding as a powerful tool for monitoring coastal and freshwater fishes within a complex and heavily urbanised ecosystem. The resulting data represents the most comprehensive fish species inventory compiled to date for the Mersey Estuary. The ecological importance of this system is underscored not only by the remarkable diversity of species detected but also by the presence of returning migratory species that depend on this ecosystem for critical life cycle stages.

Additionally, detecting species of conservation concern listed in the UK Biodiversity Action Plan (Joint Nature Conservation Committee, 2007) provides invaluable insights to support targeted conservation efforts for these vulnerable populations. However, the study also highlights a recurring pattern of unexpected species detections, such as freshwater species detected at marine sampling sites. This recurring phenomenon in tidal and lotic ecosystems underscores the complexity of distinguishing between species presence and eDNA particle distributions, as well as highlighting complex sampling dynamics in such environments. Looking ahead, integrating advanced techniques like hydrodynamic modelling and particle tracking to examine these anomalous detections will enhance data interpretation and improve the precision of species monitoring using eDNA for effective management and conservation strategies.

## Acknowledgements

This project was funded by the Mersey Gateway Environmental Trust, the University of Salford and Bangor University. We thank the members of the Molecular Ecology Group in the University of Salford for their advice on laboratory protocols and bioinformatics.

## Author contributions

ADM, IC, JMJ and PER conceived and acquired funding for the study. JMJ carried out the eDNA sampling, laboratory work and bioinformatic analyses. JMJ, NGS, CB, IC and ADM analysed the data. JMJ and ADM prepared the original draft of the manuscript, with input from IC, NGS and CB. All other authors (AD, AW and PER) contributed to editing and discussing the manuscript.

## Data Availability Statement

The data that support the findings of this study are openly available in - JmJackman27/Mersey_eDNA_data – at https://github.com/JmJackman27/Multi-season-eDNA---Mersey-estuary

## Appendix A Additional data and results that support the findings in the main text

**Table A1.**
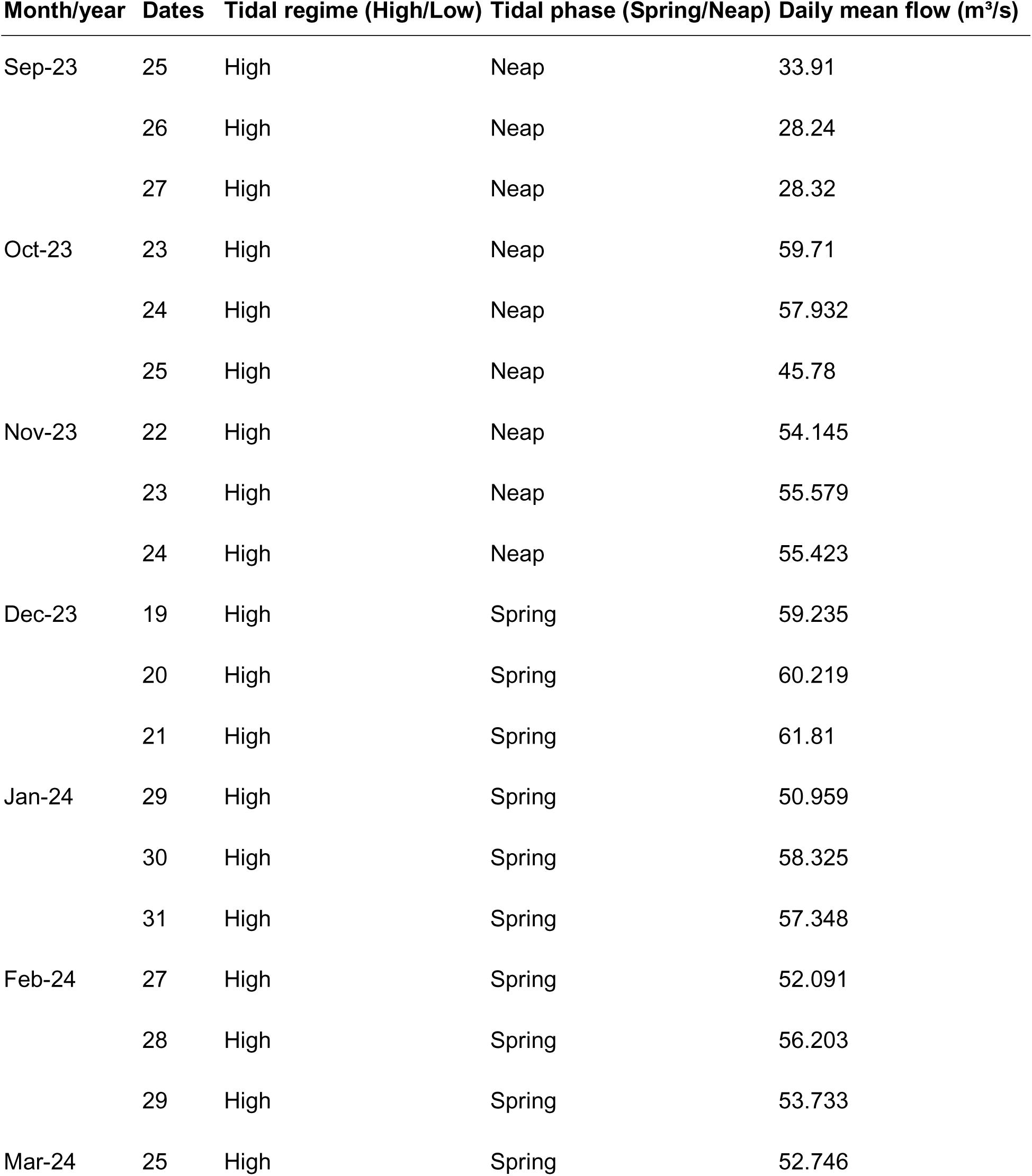
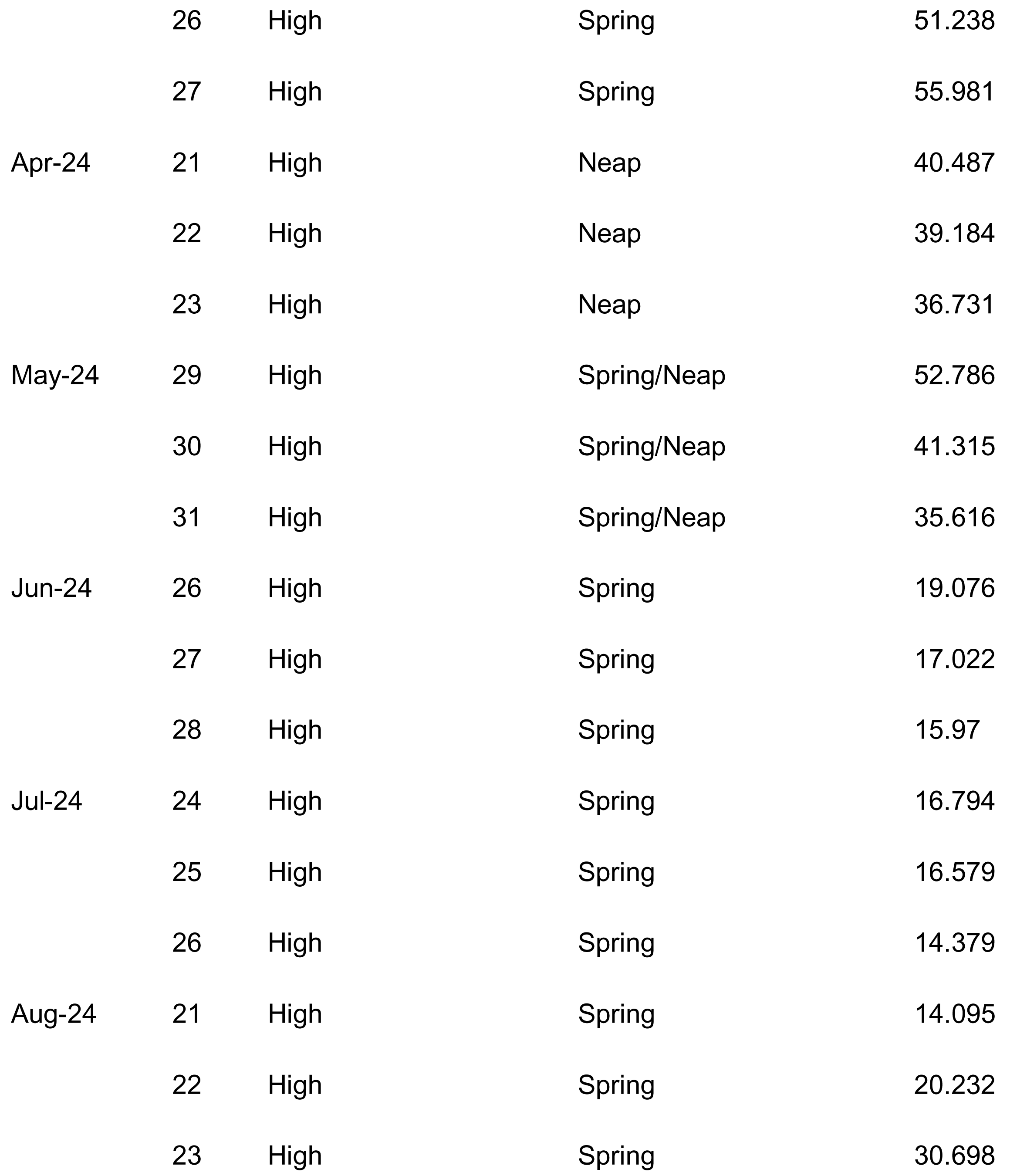
The month, year, date, tidal regime (High/Low tide), tidal phase (Spring/Neap tide) and daily average mean flow (m³/s) on each of the three sampled days during the 12 sampled months.

**Table A2.**
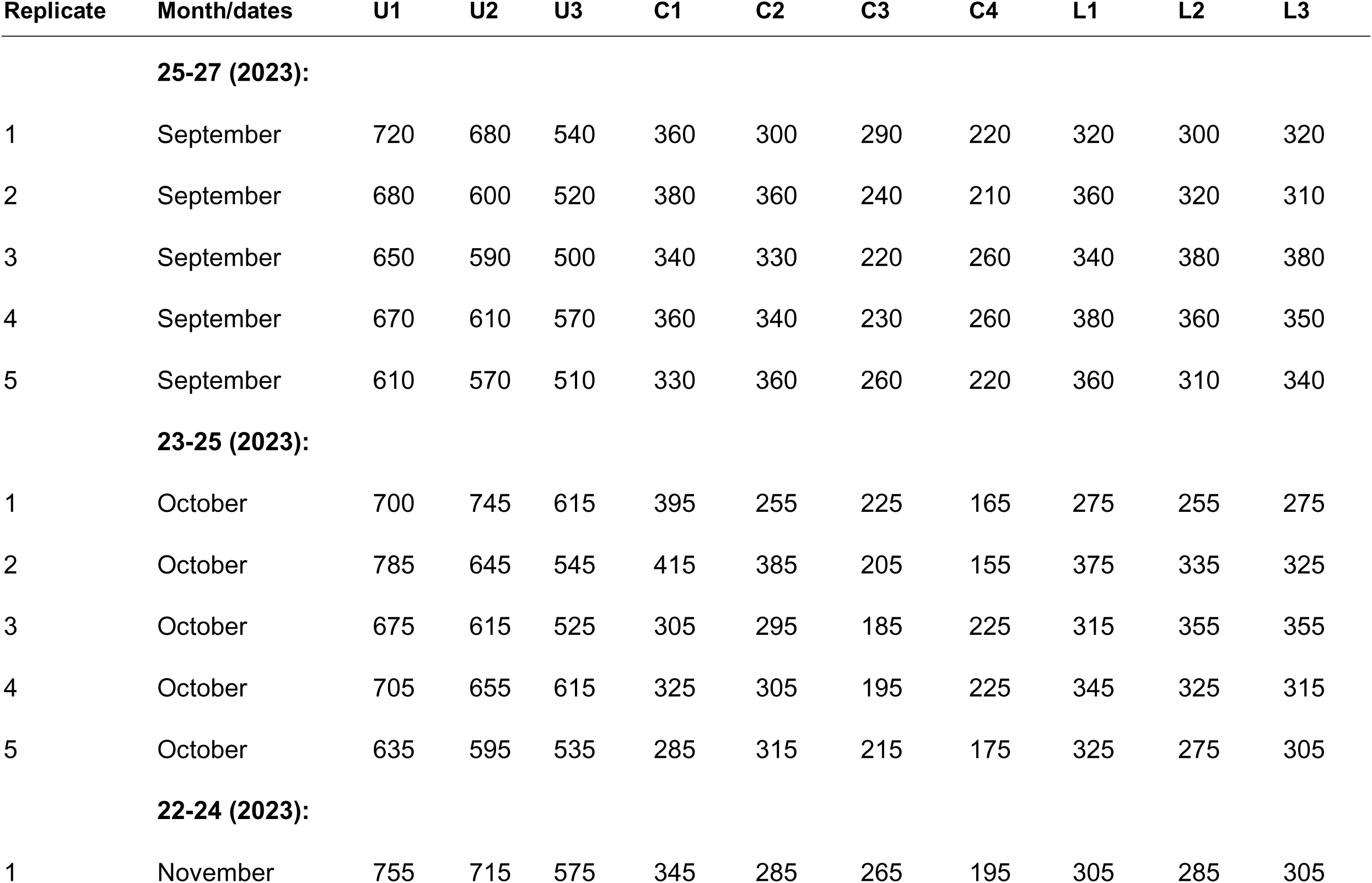
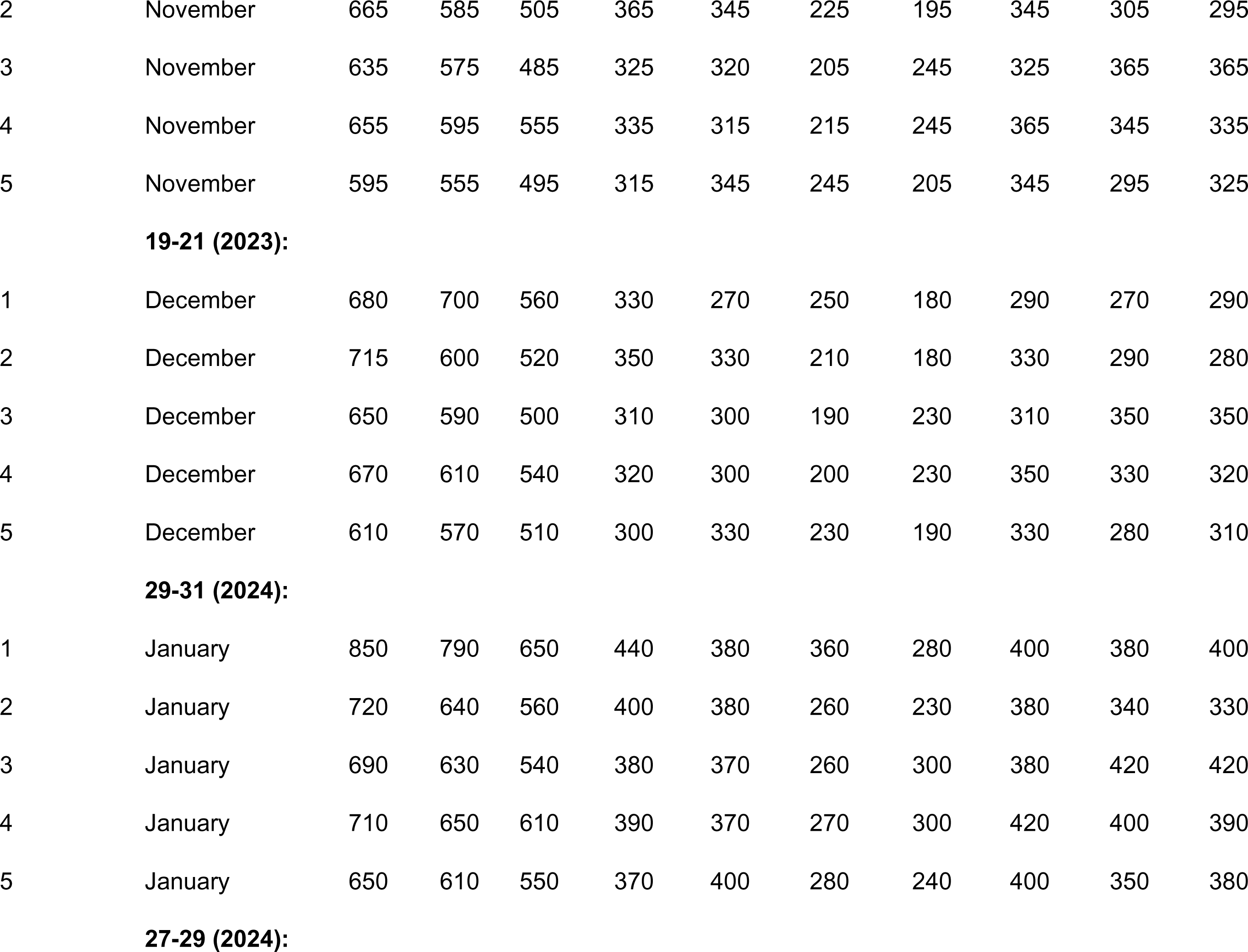
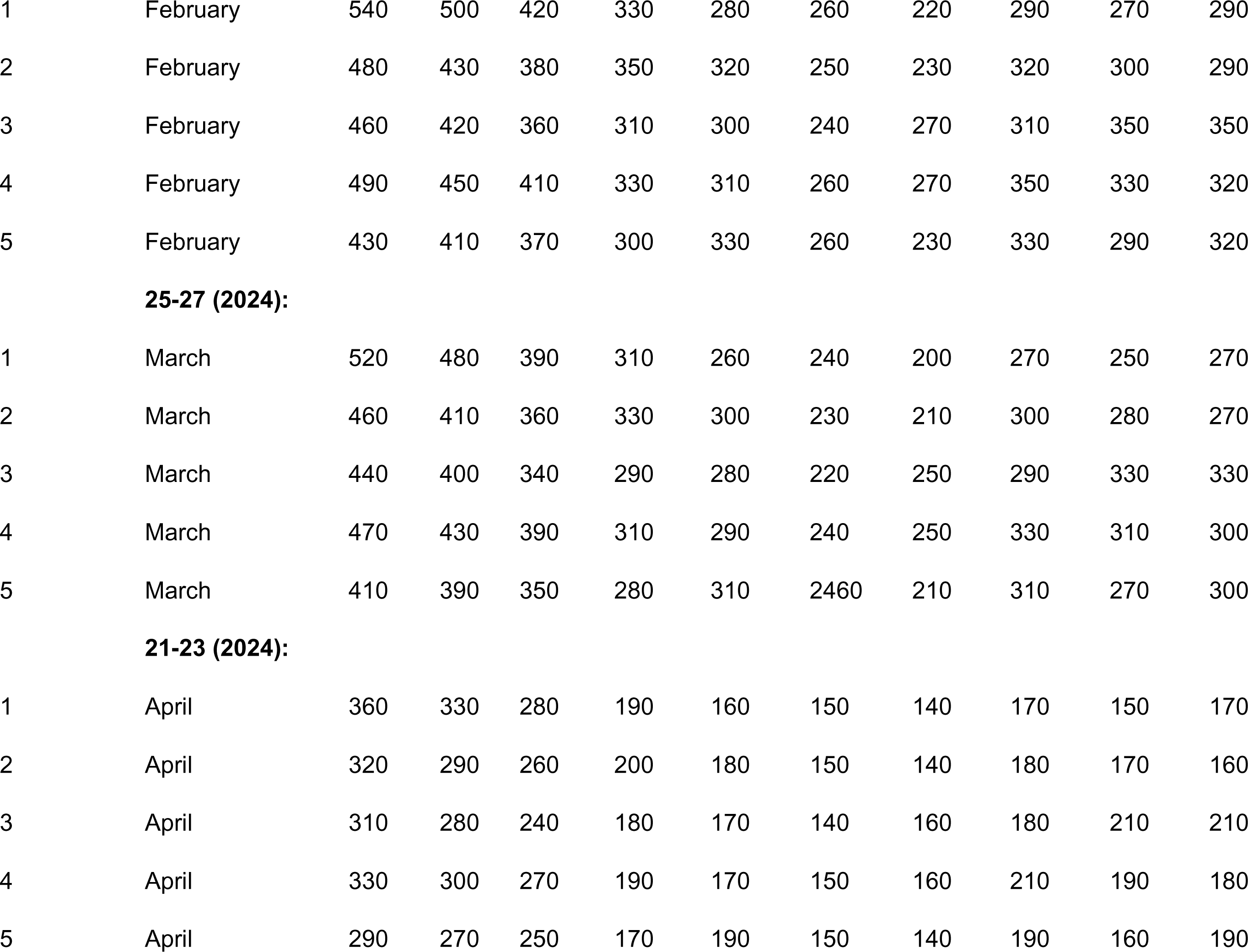
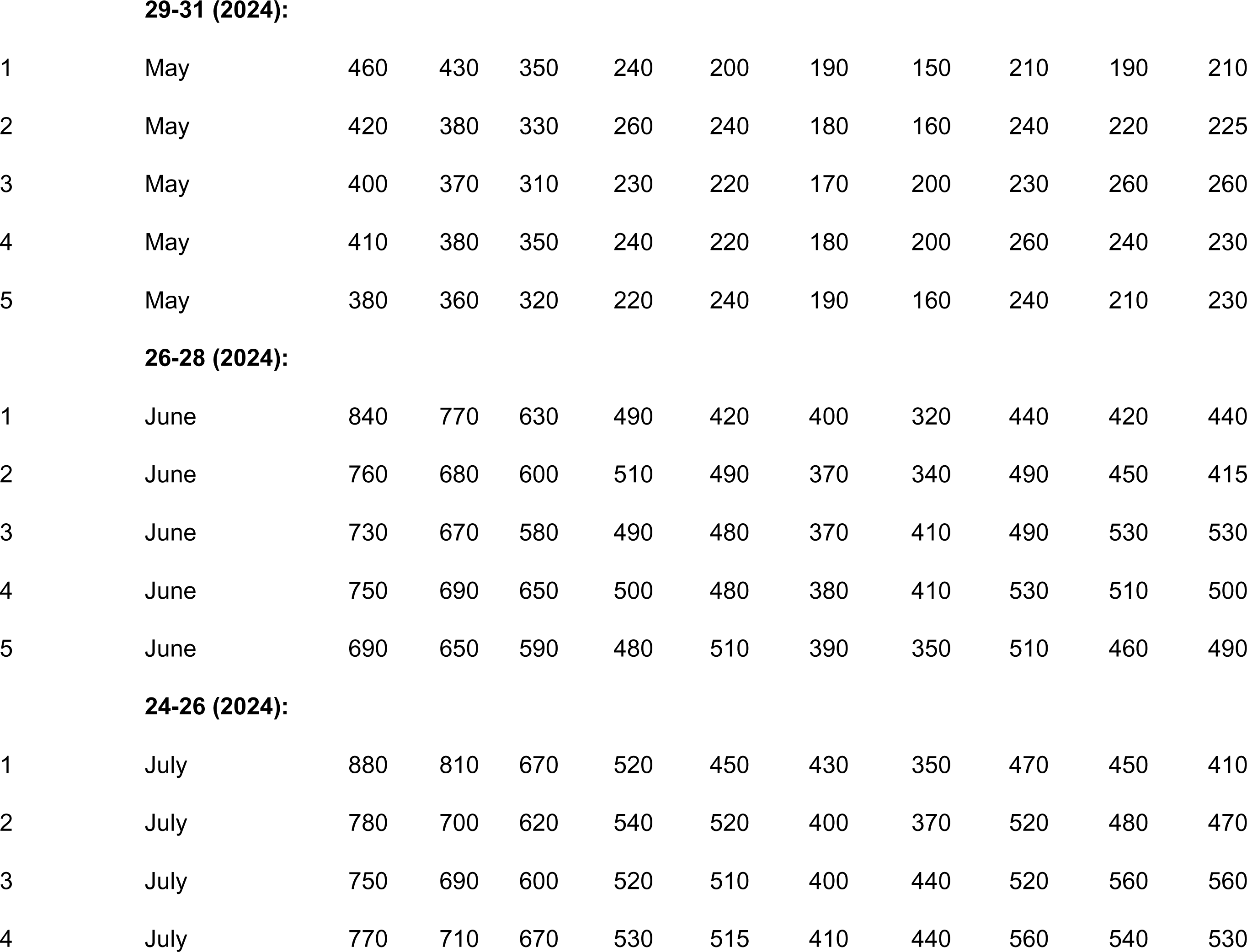
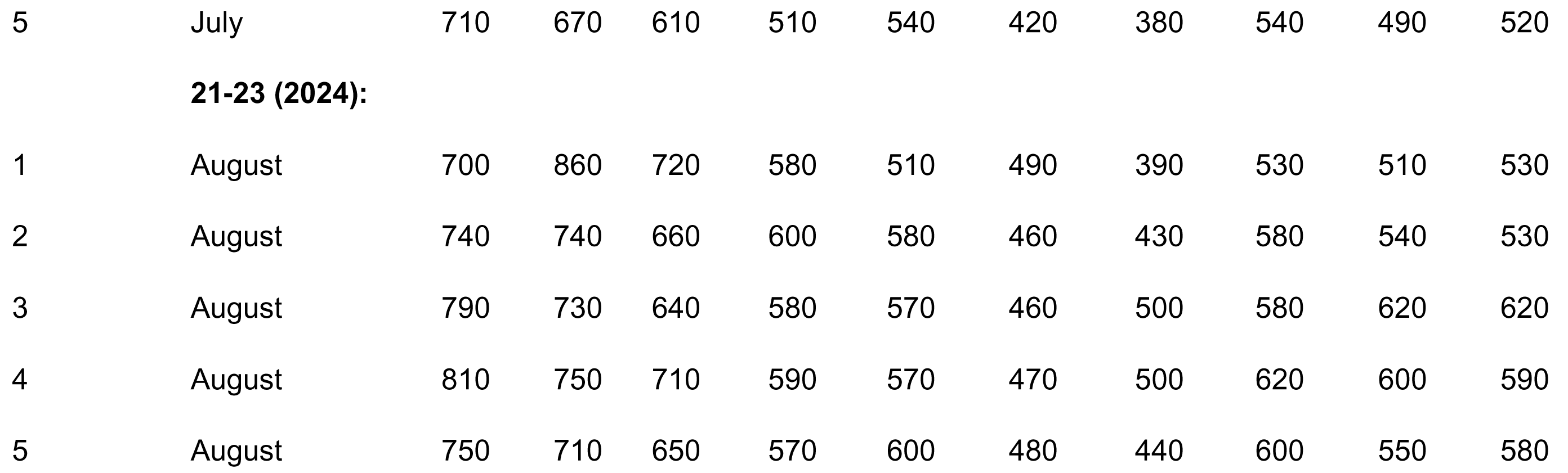
The filter (0.45 µm) sample replicates collected at each of the 10 sample locations for all 12 months and their associated dates and volumes (ml).

| Replicate | Month/dates | U1 | U2 | U3 | C1 | C2 | C3 | C4 | L1 | L2 | L3 |
| --- | --- | --- | --- | --- | --- | --- | --- | --- | --- | --- | --- |
| **25-27 (2023):** |  |  |  |  |  |  |  |  |  |  |  |
| 1 | September | 720 | 680 | 540 | 360 | 300 | 290 | 220 | 320 | 300 | 320 |
| 2 | September | 680 | 600 | 520 | 380 | 360 | 240 | 210 | 360 | 320 | 310 |
| 3 | September | 650 | 590 | 500 | 340 | 330 | 220 | 260 | 340 | 380 | 380 |
| 4 | September | 670 | 610 | 570 | 360 | 340 | 230 | 260 | 380 | 360 | 350 |
| 5 | September | 610 | 570 | 510 | 330 | 360 | 260 | 220 | 360 | 310 | 340 |
| **23-25 (2023):** |  |  |  |  |  |  |  |  |  |  |  |
| 1 | October | 700 | 745 | 615 | 395 | 255 | 225 | 165 | 275 | 255 | 275 |
| 2 | October | 785 | 645 | 545 | 415 | 385 | 205 | 155 | 375 | 335 | 325 |
| 3 | October | 675 | 615 | 525 | 305 | 295 | 185 | 225 | 315 | 355 | 355 |
| 4 | October | 705 | 655 | 615 | 325 | 305 | 195 | 225 | 345 | 325 | 315 |
| 5 | October | 635 | 595 | 535 | 285 | 315 | 215 | 175 | 325 | 275 | 305 |
| **22-24 (2023):** |  |  |  |  |  |  |  |  |  |  |  |
| 1 | November | 755 | 715 | 575 | 345 | 285 | 265 | 195 | 305 | 285 | 305 |
| 2 | November | 665 | 585 | 505 | 365 | 345 | 225 | 195 | 345 | 305 | 295 |
| 3 | November | 635 | 575 | 485 | 325 | 320 | 205 | 245 | 325 | 365 | 365 |
| 4 | November | 655 | 595 | 555 | 335 | 315 | 215 | 245 | 365 | 345 | 335 |
| 5 | November | 595 | 555 | 495 | 315 | 345 | 245 | 205 | 345 | 295 | 325 |
| **19-21 (2023):** |  |  |  |  |  |  |  |  |  |  |  |
| 1 | December | 680 | 700 | 560 | 330 | 270 | 250 | 180 | 290 | 270 | 290 |
| 2 | December | 715 | 600 | 520 | 350 | 330 | 210 | 180 | 330 | 290 | 280 |
| 3 | December | 650 | 590 | 500 | 310 | 300 | 190 | 230 | 310 | 350 | 350 |
| 4 | December | 670 | 610 | 540 | 320 | 300 | 200 | 230 | 350 | 330 | 320 |
| 5 | December | 610 | 570 | 510 | 300 | 330 | 230 | 190 | 330 | 280 | 310 |
| **29-31 (2024):** |  |  |  |  |  |  |  |  |  |  |  |
| 1 | January | 850 | 790 | 650 | 440 | 380 | 360 | 280 | 400 | 380 | 400 |
| 2 | January | 720 | 640 | 560 | 400 | 380 | 260 | 230 | 380 | 340 | 330 |
| 3 | January | 690 | 630 | 540 | 380 | 370 | 260 | 300 | 380 | 420 | 420 |
| 4 | January | 710 | 650 | 610 | 390 | 370 | 270 | 300 | 420 | 400 | 390 |
| 5 | January | 650 | 610 | 550 | 370 | 400 | 280 | 240 | 400 | 350 | 380 |
| **27-29 (2024):** |  |  |  |  |  |  |  |  |  |  |  |
| 1 | February | 540 | 500 | 420 | 330 | 280 | 260 | 220 | 290 | 270 | 290 |
| 2 | February | 480 | 430 | 380 | 350 | 320 | 250 | 230 | 320 | 300 | 290 |
| 3 | February | 460 | 420 | 360 | 310 | 300 | 240 | 270 | 310 | 350 | 350 |
| 4 | February | 490 | 450 | 410 | 330 | 310 | 260 | 270 | 350 | 330 | 320 |
| 5 | February | 430 | 410 | 370 | 300 | 330 | 260 | 230 | 330 | 290 | 320 |
| **25-27 (2024):** |  |  |  |  |  |  |  |  |  |  |  |
| 1 | March | 520 | 480 | 390 | 310 | 260 | 240 | 200 | 270 | 250 | 270 |
| 2 | March | 460 | 410 | 360 | 330 | 300 | 230 | 210 | 300 | 280 | 270 |
| 3 | March | 440 | 400 | 340 | 290 | 280 | 220 | 250 | 290 | 330 | 330 |
| 4 | March | 470 | 430 | 390 | 310 | 290 | 240 | 250 | 330 | 310 | 300 |
| 5 | March | 410 | 390 | 350 | 280 | 310 | 2460 | 210 | 310 | 270 | 300 |
| **21-23 (2024):** |  |  |  |  |  |  |  |  |  |  |  |
| 1 | April | 360 | 330 | 280 | 190 | 160 | 150 | 140 | 170 | 150 | 170 |
| 2 | April | 320 | 290 | 260 | 200 | 180 | 150 | 140 | 180 | 170 | 160 |
| 3 | April | 310 | 280 | 240 | 180 | 170 | 140 | 160 | 180 | 210 | 210 |
| 4 | April | 330 | 300 | 270 | 190 | 170 | 150 | 160 | 210 | 190 | 180 |
| 5 | April | 290 | 270 | 250 | 170 | 190 | 150 | 140 | 190 | 160 | 190 |
| **29-31 (2024):** |  |  |  |  |  |  |  |  |  |  |  |
| 1 | May | 460 | 430 | 350 | 240 | 200 | 190 | 150 | 210 | 190 | 210 |
| 2 | May | 420 | 380 | 330 | 260 | 240 | 180 | 160 | 240 | 220 | 225 |
| 3 | May | 400 | 370 | 310 | 230 | 220 | 170 | 200 | 230 | 260 | 260 |
| 4 | May | 410 | 380 | 350 | 240 | 220 | 180 | 200 | 260 | 240 | 230 |
| 5 | May | 380 | 360 | 320 | 220 | 240 | 190 | 160 | 240 | 210 | 230 |
| **26-28 (2024):** |  |  |  |  |  |  |  |  |  |  |  |
| 1 | June | 840 | 770 | 630 | 490 | 420 | 400 | 320 | 440 | 420 | 440 |
| 2 | June | 760 | 680 | 600 | 510 | 490 | 370 | 340 | 490 | 450 | 415 |
| 3 | June | 730 | 670 | 580 | 490 | 480 | 370 | 410 | 490 | 530 | 530 |
| 4 | June | 750 | 690 | 650 | 500 | 480 | 380 | 410 | 530 | 510 | 500 |
| 5 | June | 690 | 650 | 590 | 480 | 510 | 390 | 350 | 510 | 460 | 490 |
| **24-26 (2024):** |  |  |  |  |  |  |  |  |  |  |  |
| 1 | July | 880 | 810 | 670 | 520 | 450 | 430 | 350 | 470 | 450 | 410 |
| 2 | July | 780 | 700 | 620 | 540 | 520 | 400 | 370 | 520 | 480 | 470 |
| 3 | July | 750 | 690 | 600 | 520 | 510 | 400 | 440 | 520 | 560 | 560 |
| 4 | July | 770 | 710 | 670 | 530 | 515 | 410 | 440 | 560 | 540 | 530 |
| 5 | July | 710 | 670 | 610 | 510 | 540 | 420 | 380 | 540 | 490 | 520 |
| **21-23 (2024):** |  |  |  |  |  |  |  |  |  |  |  |
| 1 | August | 700 | 860 | 720 | 580 | 510 | 490 | 390 | 530 | 510 | 530 |
| 2 | August | 740 | 740 | 660 | 600 | 580 | 460 | 430 | 580 | 540 | 530 |
| 3 | August | 790 | 730 | 640 | 580 | 570 | 460 | 500 | 580 | 620 | 620 |
| 4 | August | 810 | 750 | 710 | 590 | 570 | 470 | 500 | 620 | 600 | 590 |
| 5 | August | 750 | 710 | 650 | 570 | 600 | 480 | 440 | 600 | 550 | 580 |

**Table A3.**
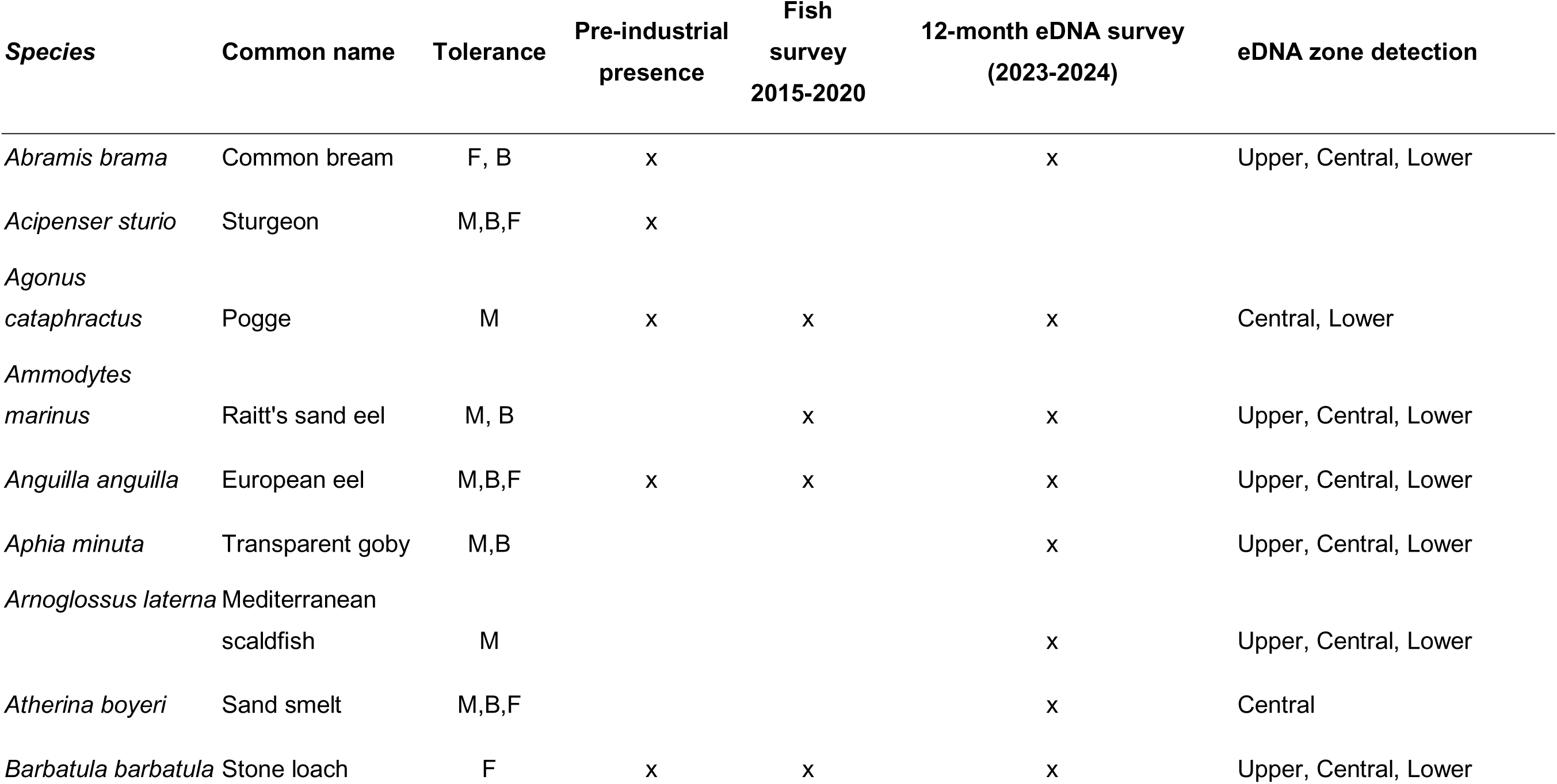
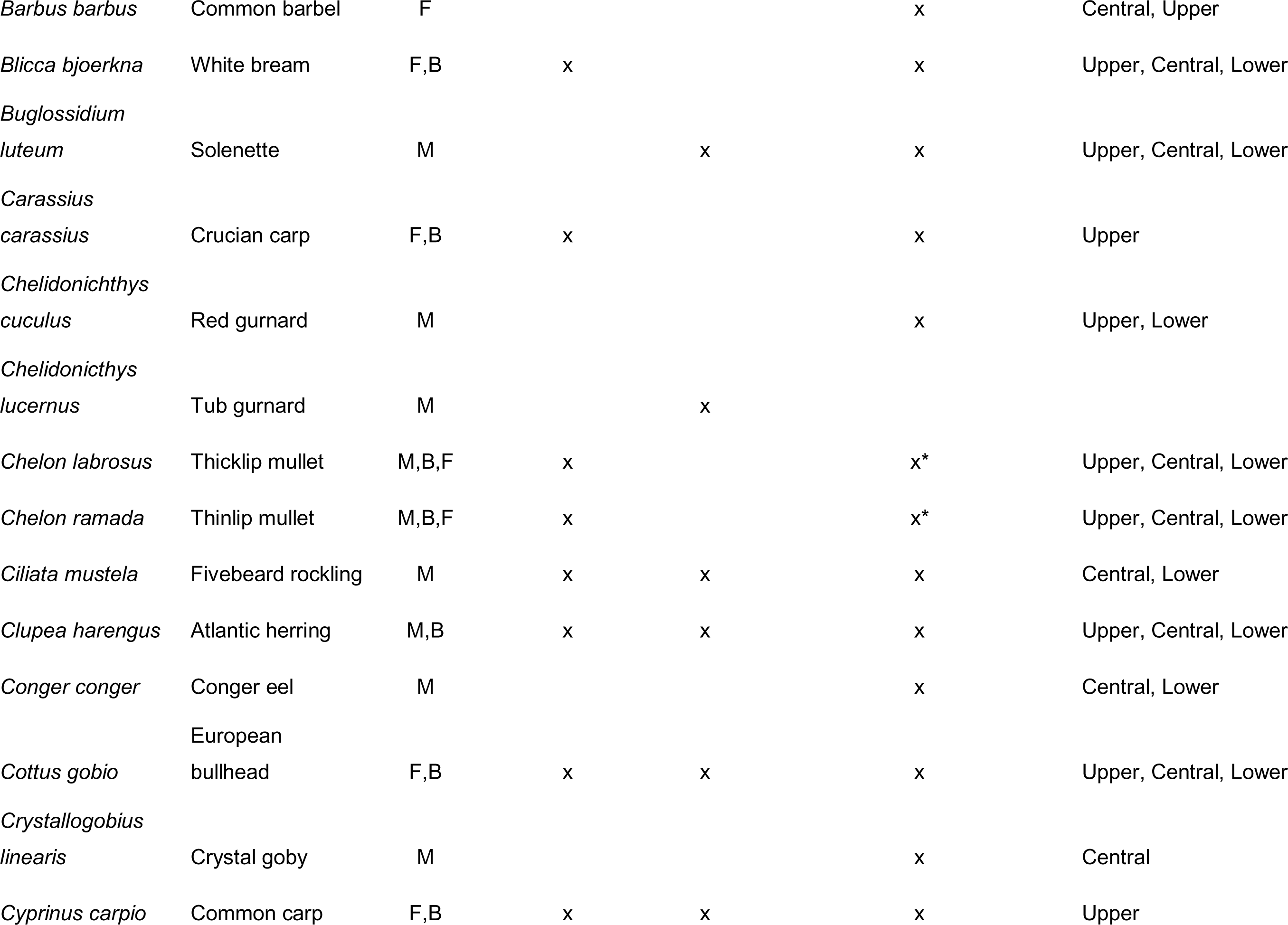
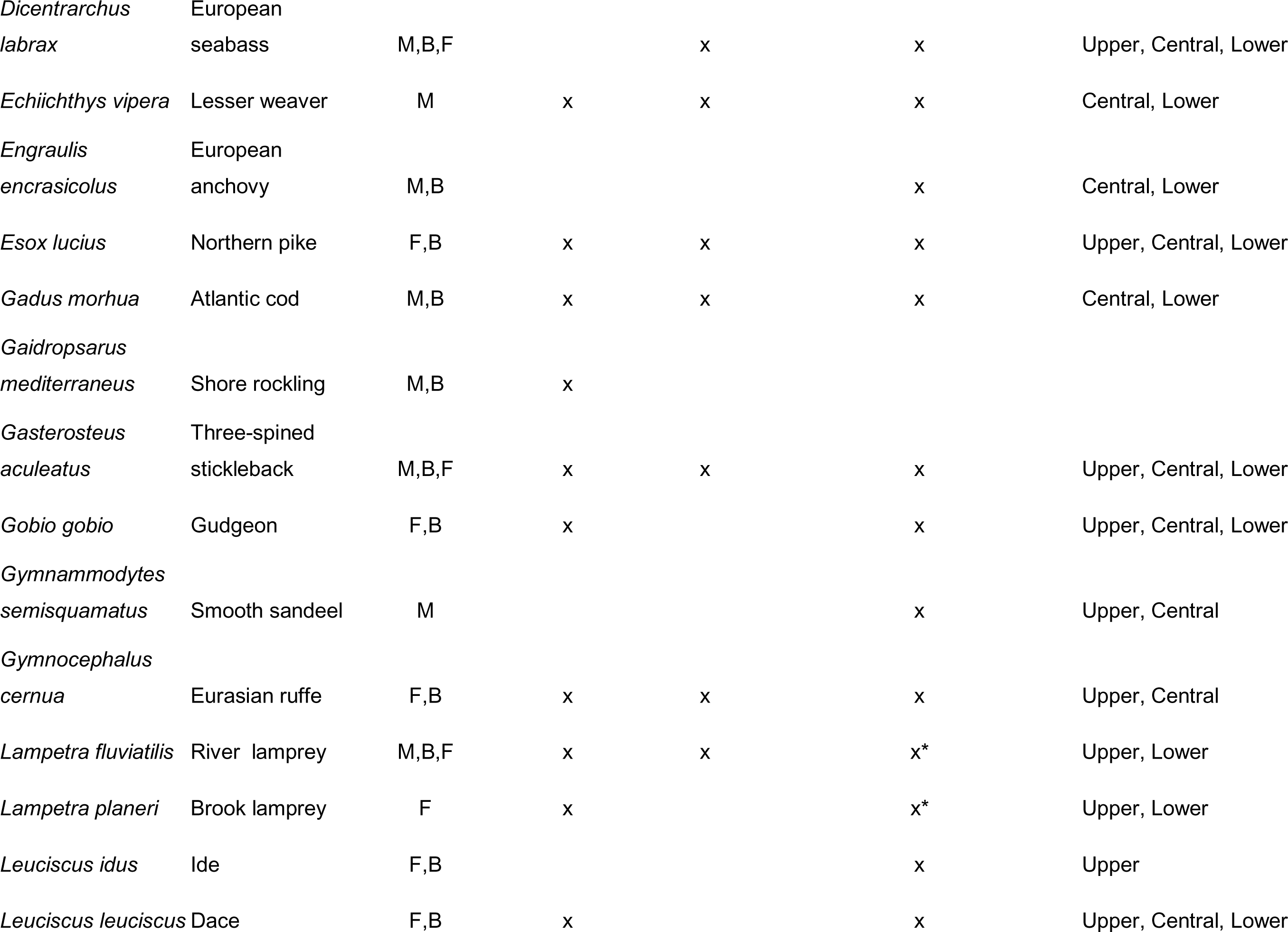
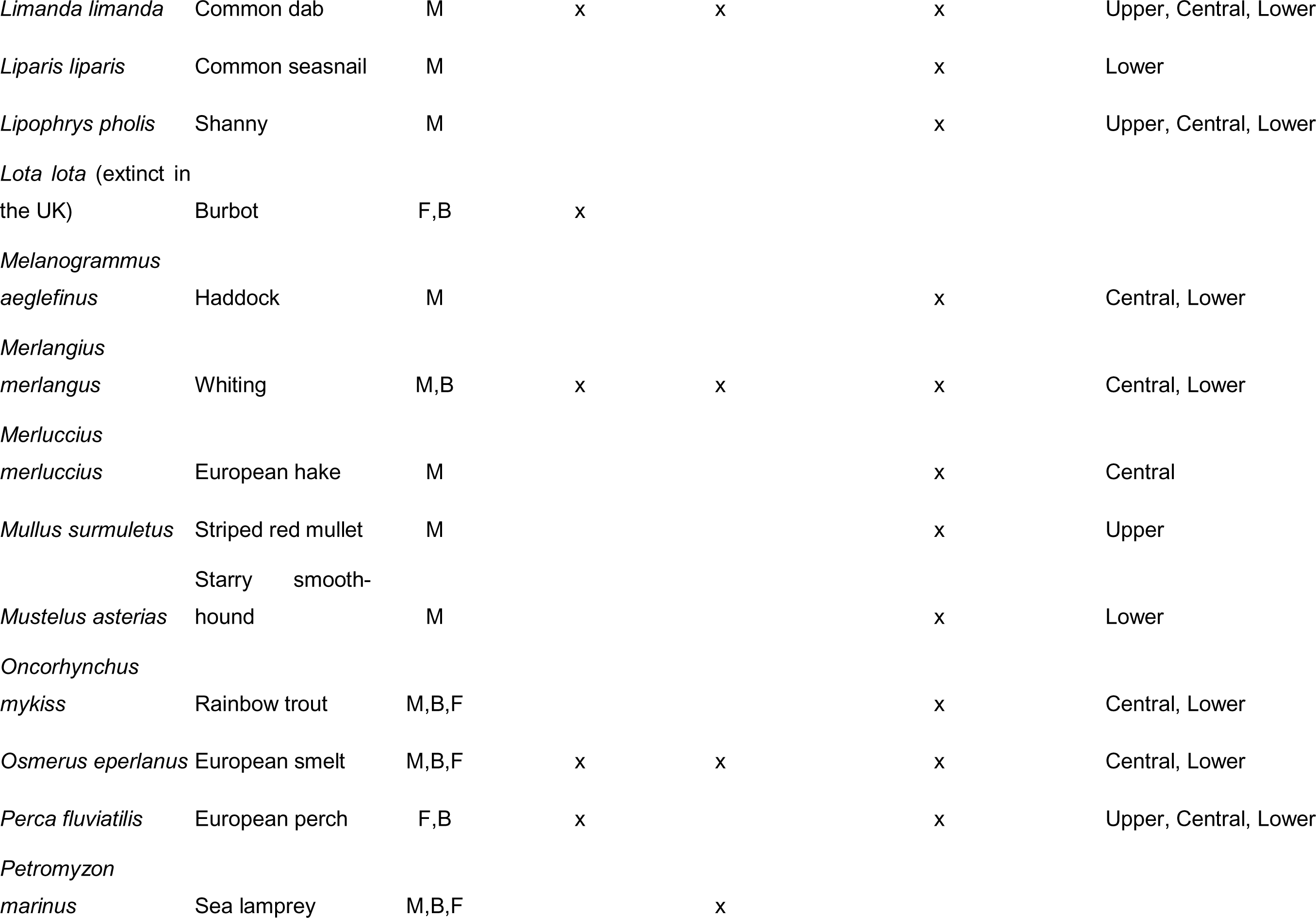
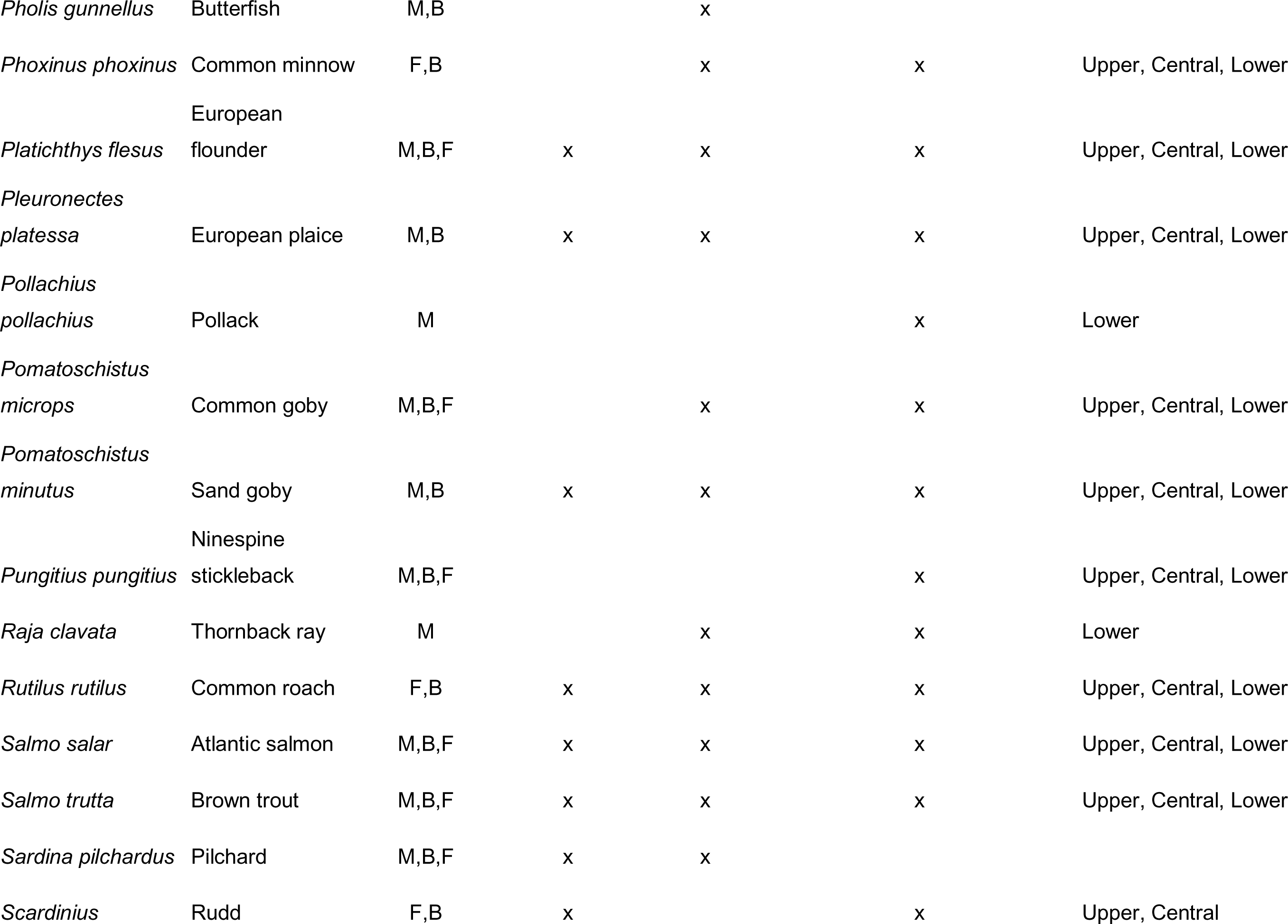
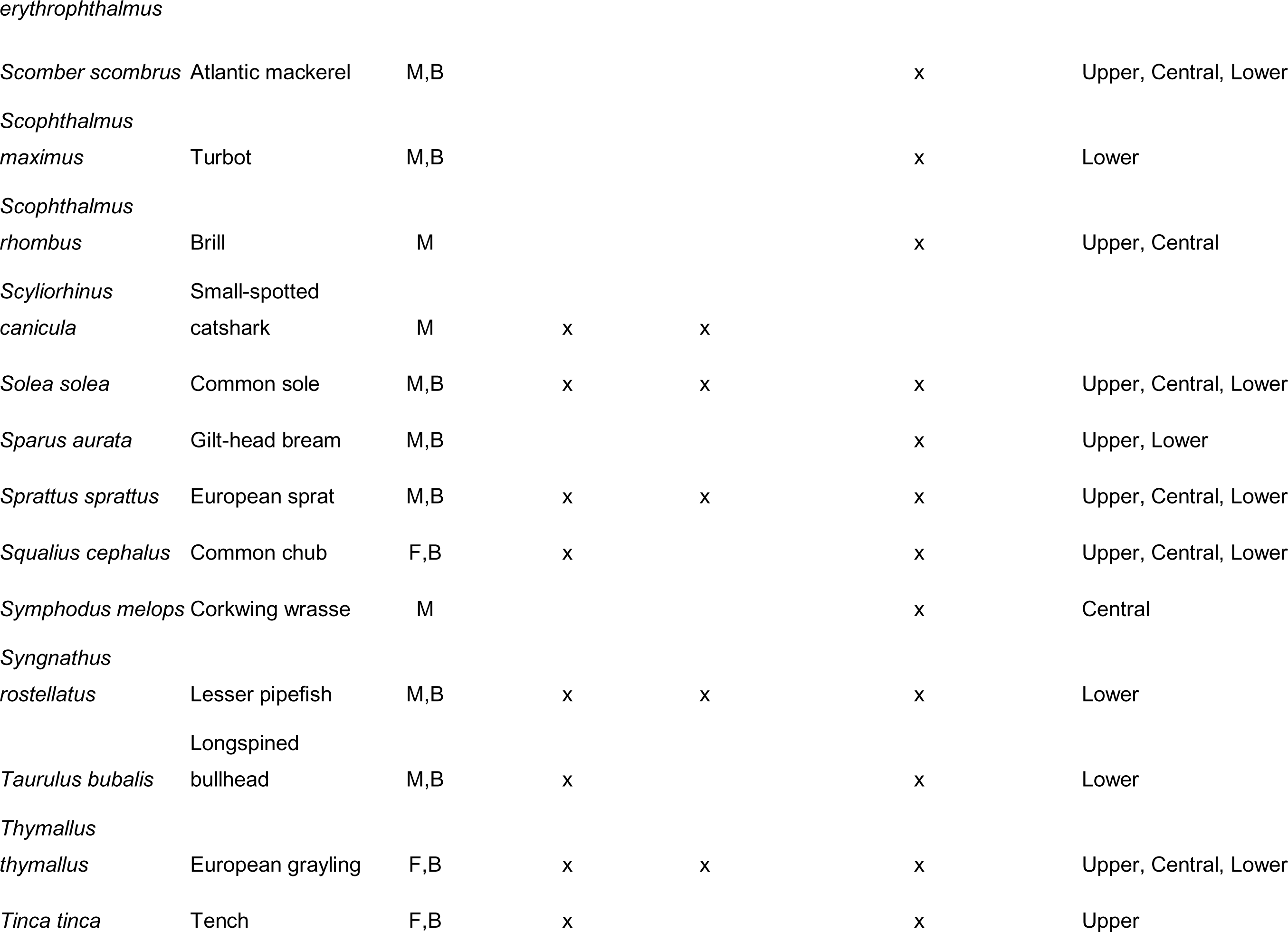
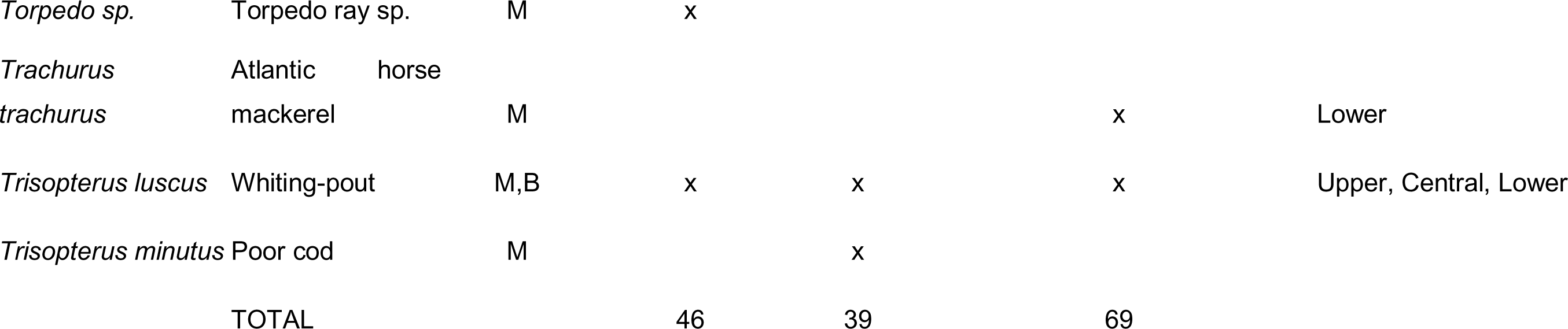
Complete list of estimated pre-industrialisation species by the Mersey Rivers Trust report (2021), the number of species identified during a five-year survey from 2015-2020, the total species eDNA detections from this study and the estuarine zones in which the eDNA for each species was detected. Each species has been classified according to its physiological saline tolerance (F = freshwater, M = marine and B = brackish). Species detected with eDNA featuring * could not be distinguished at species level.

**Table A4.** All Dunn’s post hoc comparisons of the differences in species richness between seasons. * represents a significant difference.

| **Pairs** | **Z** | **Unadjusted p value** | **Adjusted p value** |
| --- | --- | --- | --- |
| Spring vs Summer | 5.91 | 3.44e-09 | 5.16e-09* |
| Spring vs Autumn | -6.48 | 9.43e-11 | 2.83e-10* |
| Spring vs Winter | -0.51 | 0.607 | 0.607 |
| Summer vs Autumn | -0.60 | 0.552 | 0.662 |
| Summer vs Winter | -6.40 | 1.54e-10 | 3.08e-10* |
| Autumn vs Winter | -6.96 | 3.31e-12 | 1.98e-11* |

**Table A5.** All post hoc pairwise comparisons of species composition changes between sampled zones within each of the 12 months. Zones are classified as: U = Upper, C = Central and L = Lower.

| Month | Zones | Df | SumsOfSqs | F.Model | R2 | p.value | p.adjusted | sig |
| --- | --- | --- | --- | --- | --- | --- | --- | --- |
| September | U vs C | 1 | 1.65675286 | 5.7350397 | 0.15611906 | 0.001 | 0.003 | * |
|  | U vs L | 1 | 1.97284422 | 6.63964802 | 0.19737567 | 0.001 | 0.003 | * |
|  | C vs L | 1 | 0.37490888 | 0.93619001 | 0.03026197 | 0.524 | 1 |  |
| October | U vs C | 1 | 1.71738151 | 9.0970035 | 0.21609622 | 0.001 | 0.003 | * |
|  | U vs L | 1 | 2.39577592 | 15.791474 | 0.36060613 | 0.001 | 0.003 | * |
|  | C vs L | 1 | 0.80566548 | 3.93192705 | 0.10646417 | 0.002 | 0.006 | * |
| November | U vs C | 1 | 2.19077165 | 10.148758 | 0.24663583 | 0.001 | 0.003 | * |
|  | U vs L | 1 | 3.94094917 | 25.9752734 | 0.48124394 | 0.001 | 0.003 | * |
|  | C vs L | 1 | 0.73670306 | 2.65358744 | 0.07885006 | 0.013 | 0.039 | . |
| December | U vs C | 1 | 3.03254091 | 22.8169963 | 0.40878223 | 0.001 | 0.003 | * |
|  | U vs L | 1 | 4.27034186 | 36.4068716 | 0.57417864 | 0.001 | 0.003 | * |
|  | C vs L | 1 | 1.10872114 | 4.96580007 | 0.13433498 | 0.001 | 0.003 | * |
| January | U vs C | 1 | 2.40786813 | 20.8372063 | 0.39436616 | 0.001 | 0.003 | * |
|  | U vs L | 1 | 5.4740909 | 77.9199243 | 0.73564936 | 0.001 | 0.003 | * |
|  | C vs L | 1 | 1.66273065 | 10.9393087 | 0.25476211 | 0.001 | 0.003 | * |
| February | U vs C | 1 | 0.60264824 | 4.83785785 | 0.1278576 | 0.001 | 0.003 | * |
|  | U vs L | 1 | 3.74254915 | 24.387694 | 0.46552333 | 0.001 | 0.003 | * |
|  | C vs L | 1 | 2.66900281 | 13.2915901 | 0.28712753 | 0.001 | 0.003 | * |
| March | U vs C | 1 | 1.73046179 | 12.1463521 | 0.26904393 | 0.001 | 0.003 | * |
|  | U vs L | 1 | 2.82888036 | 22.6249802 | 0.44691336 | 0.001 | 0.003 | * |
|  | C vs L | 1 | 0.98920574 | 4.60337217 | 0.12241913 | 0.001 | 0.003 | * |
| April | U vs C | 1 | 1.87867947 | 16.967036 | 0.33956459 | 0.001 | 0.003 | * |
|  | U vs L | 1 | 2.66810385 | 18.5014422 | 0.39786814 | 0.001 | 0.003 | * |
|  | C vs L | 1 | 0.96138948 | 5.20161953 | 0.13616228 | 0.001 | 0.003 | * |
| May | U vs C | 1 | 2.29652745 | 22.5405163 | 0.40583916 | 0.001 | 0.003 | * |
|  | U vs L | 1 | 2.0777841 | 11.8021264 | 0.29652 | 0.001 | 0.003 | * |
|  | C vs L | 1 | 0.81467615 | 4.35377119 | 0.11655506 | 0.001 | 0.003 | * |
| June | U vs C | 1 | 2.35447627 | 16.0367319 | 0.32703509 | 0.001 | 0.003 | * |
|  | U vs L | 1 | 1.63107235 | 8.49351408 | 0.23274037 | 0.001 | 0.003 | * |
|  | C vs L | 1 | 0.77417728 | 3.76225983 | 0.10234028 | 0.001 | 0.003 | * |
| July | U vs C | 1 | 1.48839175 | 5.89720123 | 0.15561047 | 0.001 | 0.003 | * |
|  | U vs L | 1 | 2.15534262 | 9.29555698 | 0.24924033 | 0.001 | 0.003 | * |
|  | C vs L | 1 | 0.29677638 | 0.85928381 | 0.02615041 | 0.54 | 1 |  |
| August | U vs C | 1 | 1.29007957 | 8.6838875 | 0.2083272 | 0.001 | 0.003 | * |
|  | U vs L | 1 | 0.81154339 | 4.62251237 | 0.1461779 | 0.001 | 0.003 | * |
|  | C vs L | 1 | 0.58450753 | 3.04287146 | 0.08683282 | 0.015 | 0.045 | . |

**Table A6.** All post hoc pairwise comparisons of monthly species composition changes for each of the 12 months.

| pairs | Df | SumsOfSqs | F.Model | R2 | P value | sig |
| --- | --- | --- | --- | --- | --- | --- |
| Sep vs Oct | 1 | 85349.1782 | 1233.67098 | 0.92640825 | 0.005 | * |
| Sep vs Nov | 1 | 14768.0065 | 85.7994316 | 0.4668101 | 0.006 | * |
| Sep vs Dec | 1 | 1388.95758 | 23.0727434 | 0.19056926 | 0.071 |  |
| Sep vs Jan | 1 | -15854.842 | -47.038308 | -1.0234242 | 1 |  |
| Sep vs Feb | 1 | 1259.01663 | 19.0871459 | 0.16301658 | 0.088 |  |
| Sep vs Mar | 1 | -59232.453 | -45.754885 | -0.8757734 | 1 |  |
| Sep vs Apr | 1 | -3103.9484 | -40.435777 | -0.7024463 | 1 |  |
| Sep vs May | 1 | -2443.6666 | -22.216575 | -0.2931588 | 0.992 |  |
| Sep vs Jun | 1 | -396333.24 | -48.85031 | -0.9939088 | 1 |  |
| Sep vs Jul | 1 | -2538.988 | -34.205923 | -0.5361928 | 1 |  |
| Sep vs Aug | 1 | -168192.72 | -46.649816 | -0.9084644 | 0.998 |  |
| Oct vs Nov | 1 | -5601.0783 | -45.929891 | -0.8820779 | 1 |  |
| Oct vs Dec | 1 | 6.69365989 | 0.667692 | 0.00676708 | 0.507 |  |
| Oct vs Jan | 1 | 89515.3121 | 314.983118 | 0.77204939 | 0.002 | * |
| Oct vs Feb | 1 | -212.14108 | -13.437282 | -0.1589031 | 0.983 |  |
| Oct vs Mar | 1 | 695986.703 | 559.301203 | 0.85090549 | 0.004 | * |
| Oct vs Apr | 1 | -1121.6232 | -42.184629 | -0.7557887 | 1 |  |
| Oct vs May | 1 | 10330.1143 | 172.689638 | 0.63796176 | 0.003 | * |
| Oct vs Jun | 1 | -393858.38 | -48.847353 | -0.9937889 | 1 |  |
| Oct vs Jul | 1 | 29244.7926 | 1215.87001 | 0.92541119 | 0.004 | * |
| Oct vs Aug | 1 | -168744.99 | -47.463508 | -0.9391928 | 1 |  |
| Nov vs Dec | 1 | -4612.685 | -40.833084 | -0.7142783 | 0.998 |  |
| Nov vs Jan | 1 | -17694.998 | -45.063908 | -0.940083 | 1 |  |
| Nov vs Feb | 1 | -4651.8405 | -39.181038 | -0.6661294 | 0.999 |  |
| Nov vs Mar | 1 | -65358.265 | -48.509631 | -0.9801833 | 1 |  |
| Nov vs Apr | 1 | -3711.491 | -28.654019 | -0.4132037 | 1 |  |
| Nov vs May | 1 | -5974.7992 | -36.709646 | -0.5989465 | 1 |  |
| Nov vs Jun | 1 | -387346.21 | -47.434115 | -0.9380655 | 1 |  |
| Nov vs Jul | 1 | 31744.5326 | 249.972899 | 0.71836887 | 0.002 | * |
| Nov vs Aug | 1 | -177654.89 | -48.563514 | -0.9823415 | 1 |  |
| Dec vs Jan | 1 | -13250.864 | -48.233406 | -1.0774419 | 1 |  |
| Dec vs Feb | 1 | 16383.5331 | 2408.11514 | 0.96089565 | 0.001 | * |
| Dec vs Mar | 1 | -58091.657 | -47.022466 | -0.9224155 | 0.999 |  |
| Dec vs Apr | 1 | 5989.40918 | 340.222236 | 0.77636918 | 0.001 | * |
| Dec vs May | 1 | -2143.0633 | -42.157299 | -0.7549295 | 0.999 |  |
| Dec vs Jun | 1 | -383643.49 | -47.633549 | -0.9457396 | 0.999 |  |
| Dec vs Jul | 1 | 272.541414 | 18.0867907 | 0.15580404 | 0.028 | * |
| Dec vs Aug | 1 | -102859.37 | -29.004919 | -0.4203911 | 0.992 |  |
| Jan vs Feb | 1 | 1582342.13 | 5635.20151 | 0.98376454 | 0.002 | * |
| Jan vs Mar | 1 | -63675.344 | -40.417283 | -0.768642 | 0.999 |  |
| Jan vs Apr | 1 | 11640.5606 | 39.8406953 | 0.29991333 | 0.024 | * |
| Jan vs May | 1 | 11995061.2 | 36660.3096 | 0.99746962 | 0.004 | * |
| Jan vs Jun | 1 | -383006.38 | -43.71869 | -0.8871251 | 1 |  |
| Jan vs Jul | 1 | -8381.643 | -28.95159 | -0.4520267 | 0.995 |  |
| Jan vs Aug | 1 | -159450.75 | -39.757726 | -0.7467323 | 0.998 |  |
| Feb vs Mar | 1 | -60464.916 | -48.716275 | -0.988486 | 1 |  |
| Feb vs Apr | 1 | -62.296108 | -2.666006 | -0.0279649 | 0.692 |  |
| Feb vs May | 1 | -1573.7188 | -27.805519 | -0.3961212 | 0.997 |  |
| Feb vs Jun | 1 | -385442.16 | -47.822658 | -0.9530728 | 1 |  |
| Feb vs Jul | 1 | 45750.1708 | 2196.25826 | 0.95728467 | 0.001 | * |
| Feb vs Aug | 1 | -117599.33 | -33.107586 | -0.5101919 | 0.998 |  |
| Mar vs Apr | 1 | -60291.907 | -48.157801 | -0.9662054 | 1 |  |
| Mar vs May | 1 | -59706.659 | -46.457238 | -0.9013339 | 1 |  |
| Mar vs Jun | 1 | -449716.17 | -48.416856 | -0.9764781 | 1 |  |
| Mar vs Jul | 1 | -59778.624 | -47.844729 | -0.9539322 | 0.999 |  |
| Mar vs Aug | 1 | -233718.17 | -48.888525 | -0.9954603 | 1 |  |
| Apr vs May | 1 | -1893.7629 | -28.098085 | -0.4019644 | 0.998 |  |
| Apr vs Jun | 1 | -383142.68 | -47.473737 | -0.9395854 | 1 |  |
| Apr vs Jul | 1 | -1118.1759 | -35.349631 | -0.5642366 | 1 |  |
| Apr vs Aug | 1 | -171820.36 | -48.225717 | -0.9688882 | 0.998 |  |
| May vs Jun | 1 | -363936.84 | -44.909104 | -0.8458909 | 0.999 |  |
| May vs Jul | 1 | 131674.801 | 2030.06299 | 0.95394873 | 0.003 | * |
| May vs Aug | 1 | -174388.91 | -48.49434 | -0.9795716 | 0.999 |  |
| Jun vs Jul | 1 | -386310.43 | -47.881286 | -0.9553574 | 1 |  |
| Jun vs Aug | 1 | -567643.56 | -48.937773 | -0.9974633 | 1 |  |
| Jul vs Aug | 1 | -173387.1 | -48.700123 | -0.9878346 | 0.997 |  |

**Figure A1.**
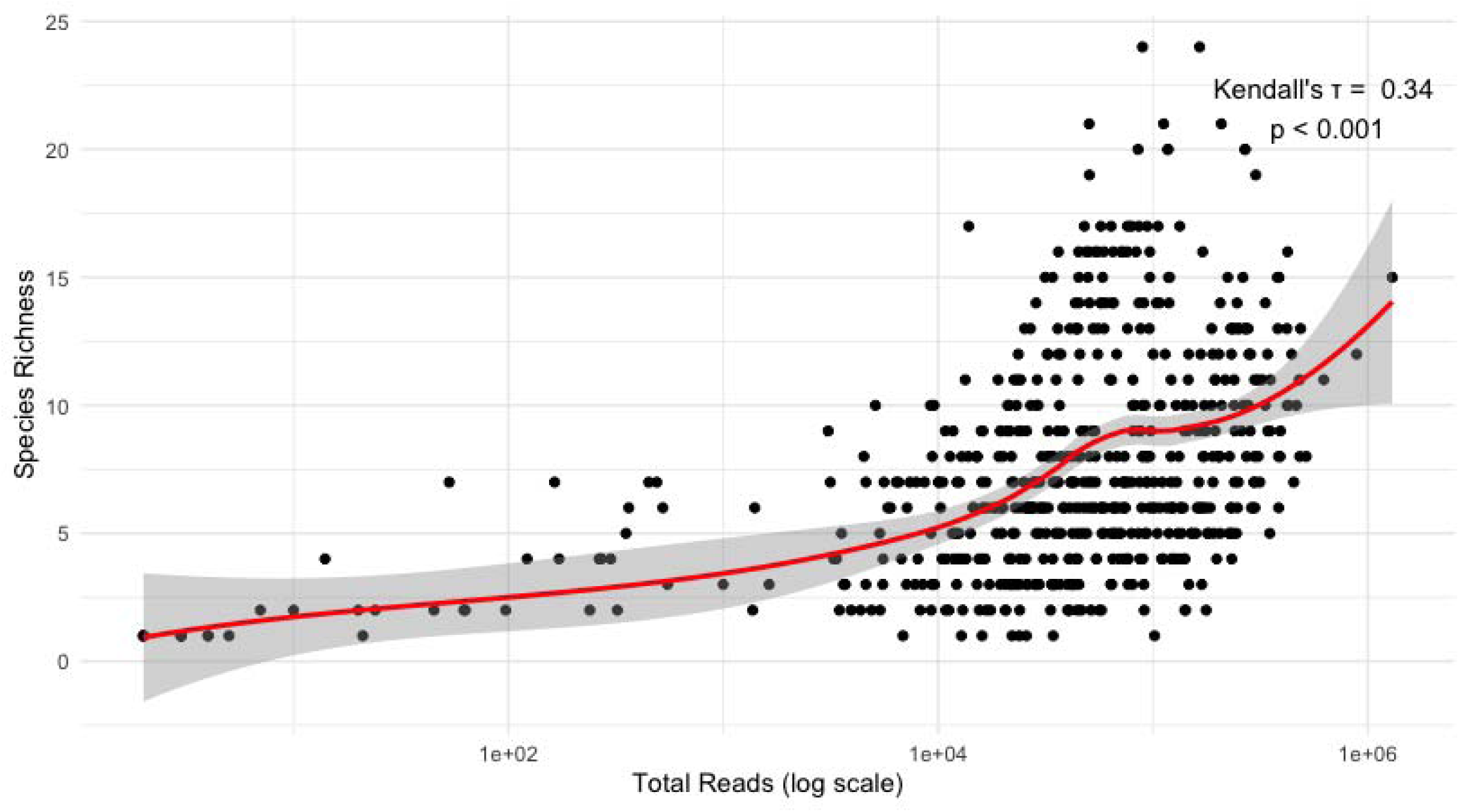
Kendall’s Tau correlation between total sequencing reads and species richness. A weak but statistically significant positive correlation (τ = 0.34, p < 0.001).

**Figure A2.**
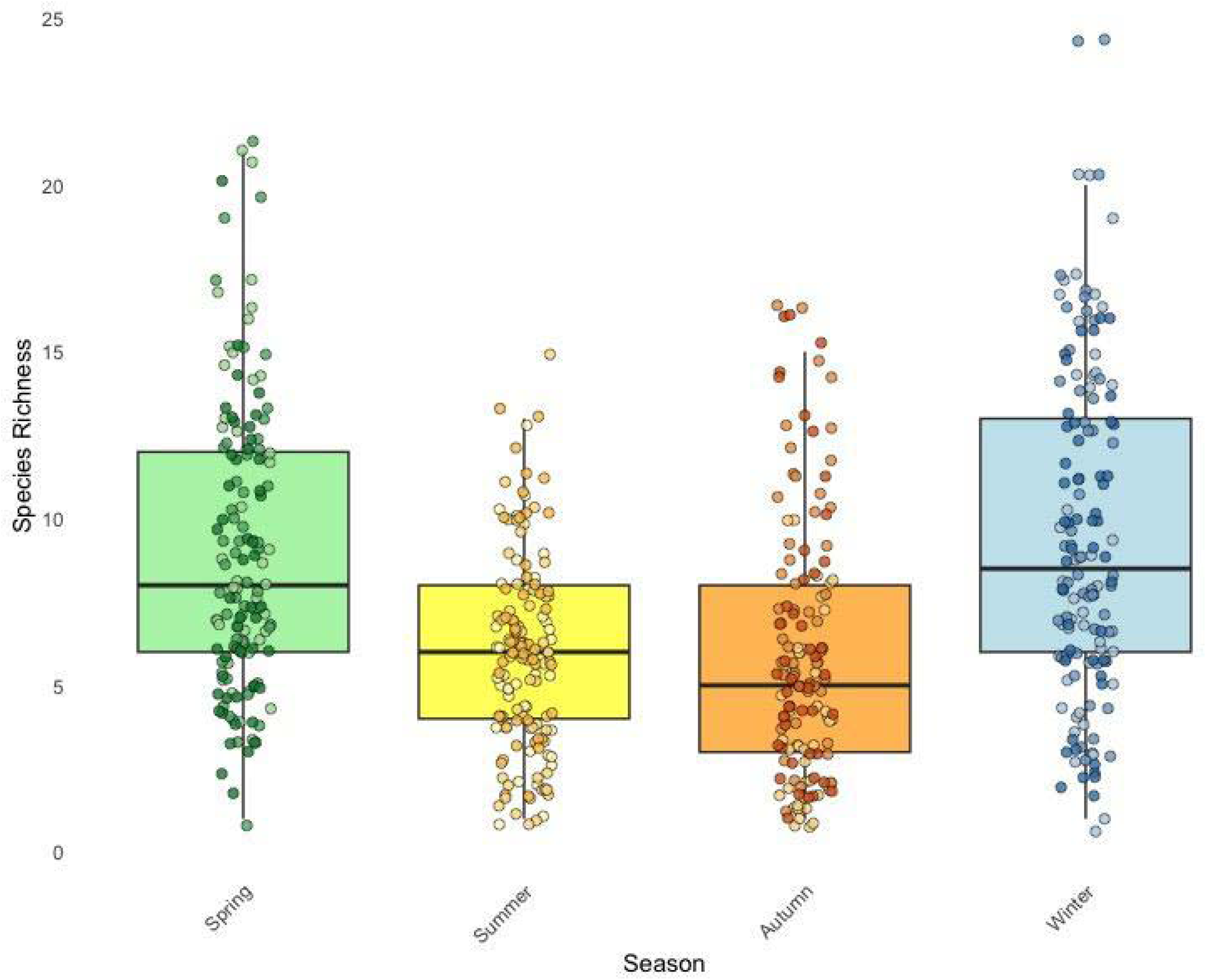
Overall species richness that was detected during each season (autumn, spring, summer and winter). The jitter points overlaid on each box represent the variation in species richness between the three months (colour gradient from darkest in month 1 to lightest in month 3 to differentiate) within each season.

**Figure A3.**
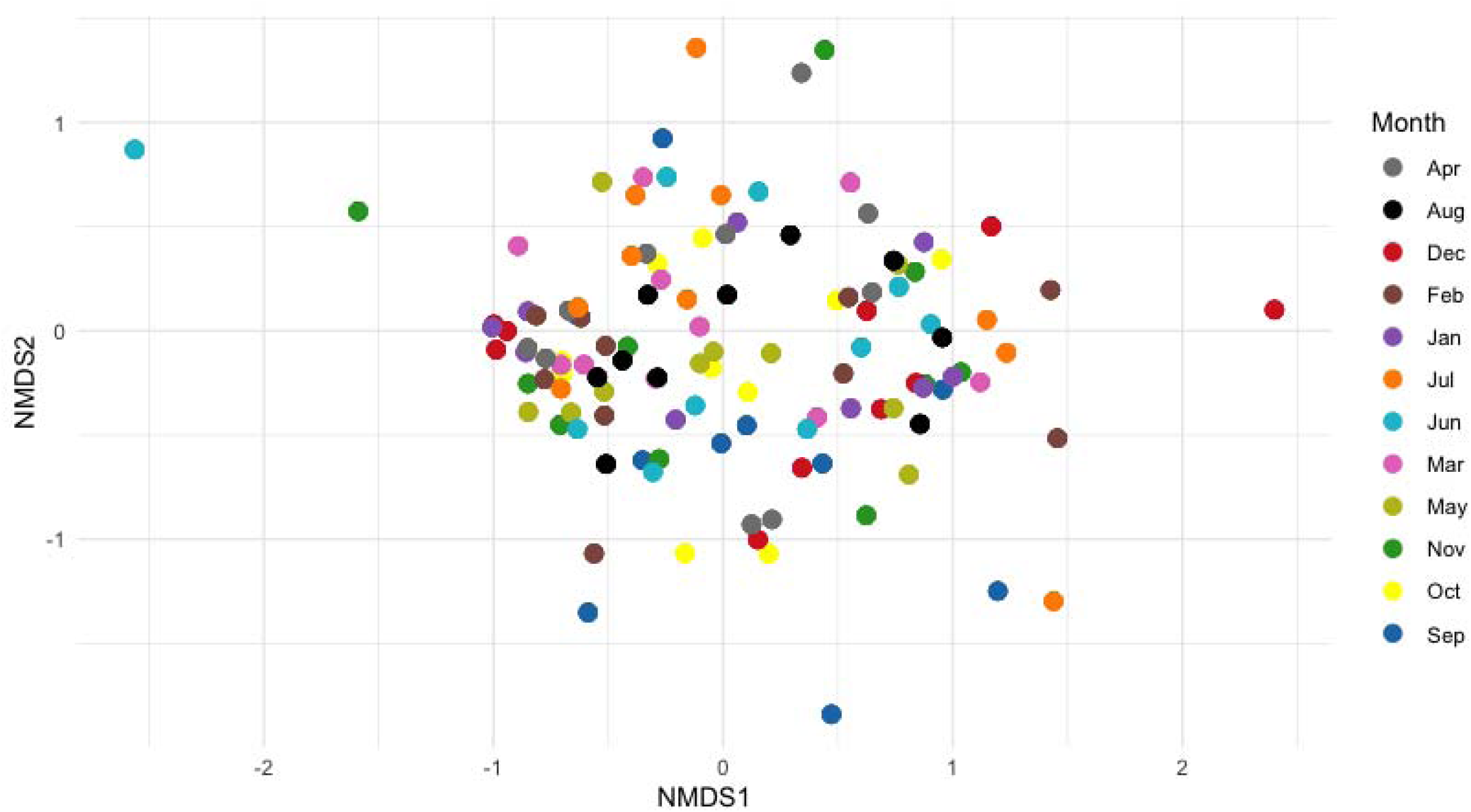
A nonmetric multidimensional scaling (NMDS) analysis using Bray–Curtis dissimilarity (stress = 0.161). The points in the plot represent the ten sampling sites for each of the 12 months sampled (10 x 12 sites = 120 points). Each month’s ten points are colour-coded to differentiate between months.

**Figure A4.**
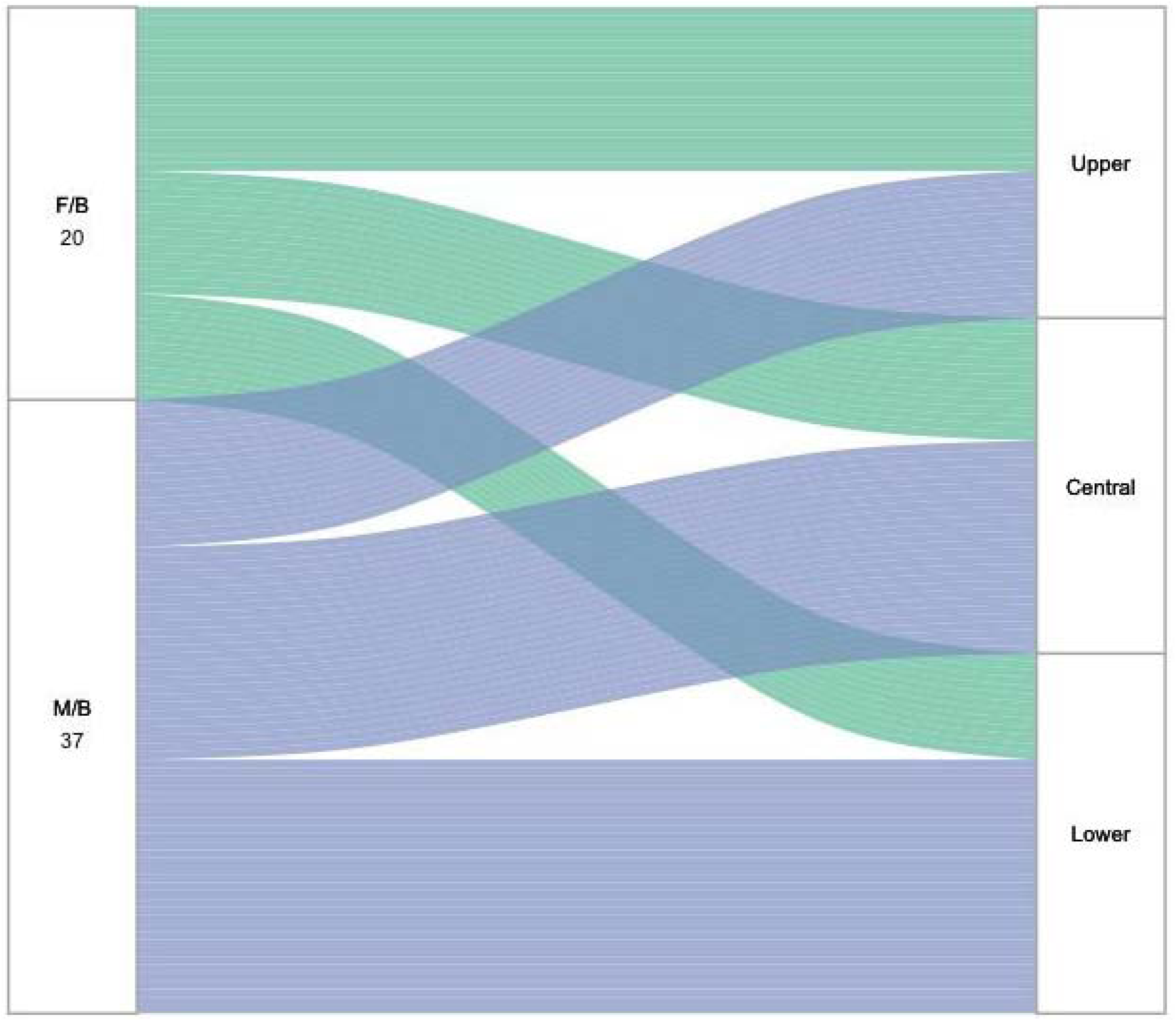
Alluvial plot showing detections of freshwater/brackish (green) and marine/brackish (blue) species (left) and different habitat zones (right) in which the eDNA signal was detected. Species classifications are based on known physiological tolerances (see Table A3), with detections outside their expected zones indicating potential eDNA transport via tidal dynamics.

